# Distinct gene expression by expanded clones of quiescent memory CD4^+^ T cells harboring intact latent HIV-1 proviruses

**DOI:** 10.1101/2022.03.02.482683

**Authors:** Georg H J Weymar, Yotam Bar-On, Thiago Y. Oliveira, Christian Gaebler, Victor Ramos, Harald Hartweger, Gaëlle Breton, Marina Caskey, Lillian B Cohn, Mila Jankovic, Michel C Nussenzweig

## Abstract

Antiretroviral therapy controls but does not cure HIV-1 infection due to a reservoir of rare CD4^+^ T cells harboring latent proviruses. Little is known about the transcriptional program of latent cells. Here we report a novel strategy to enrich clones of latent cells carrying intact, replication-competent HIV-1 proviruses from blood based on their expression of unique T cell receptors. Latent cell enrichment enabled single cell transcriptomic analysis of 1,050 CD4^+^ T cells belonging to expanded clones harboring intact HIV-1 proviruses from 6 different individuals. The analysis revealed that most of these cells are T effector memory cells that are enriched for expression of HLA-DR, HLA-DP, CD74, CCL5, Granzymes A and K, cystatin F, LYAR and DUSP2. We conclude that expanded clones of latent cells carrying intact HIV-1 proviruses persist preferentially in a distinct CD4^+^ T cell population opening new possibilities for eradication.

**Graphical Abstract:** 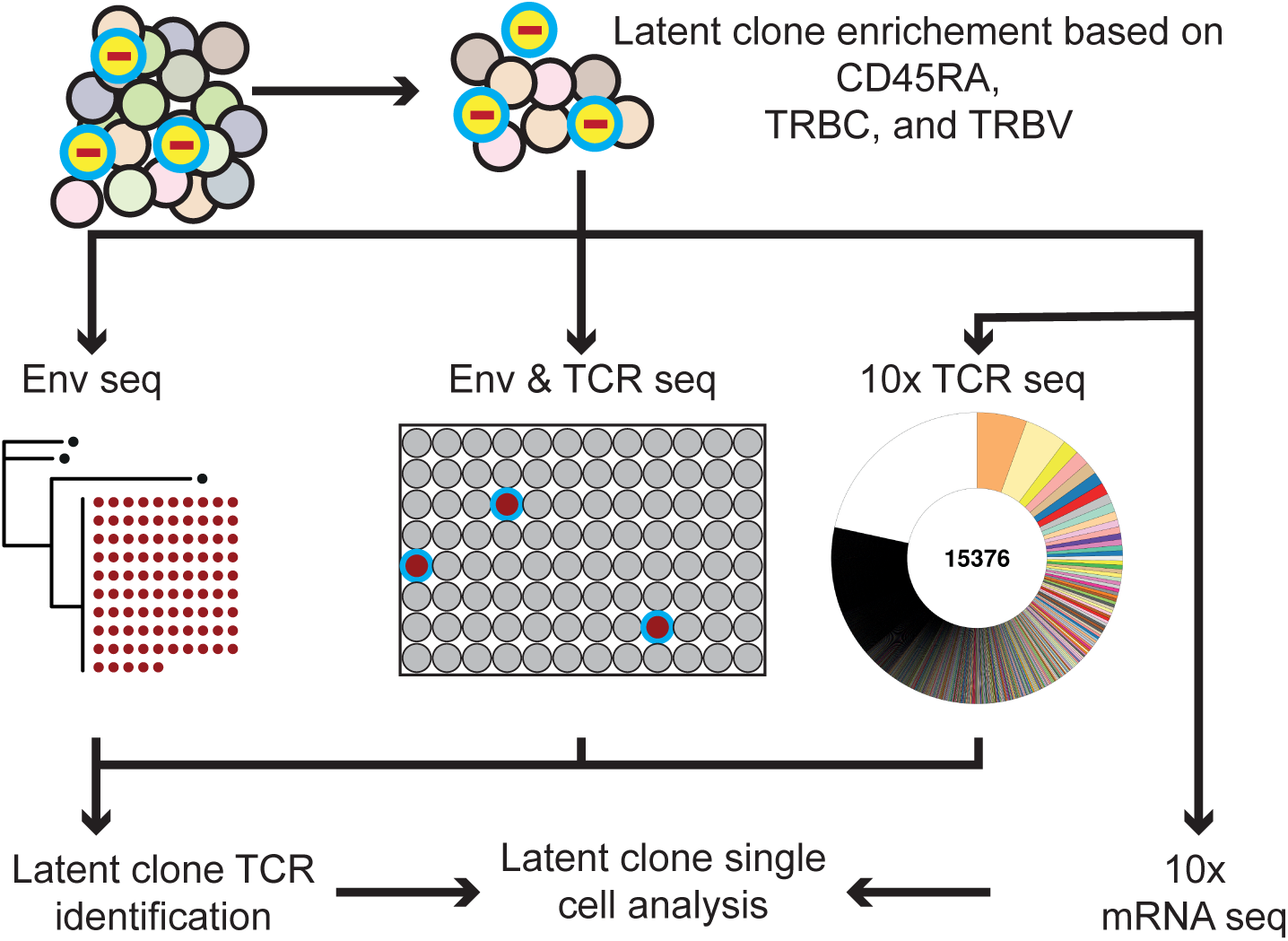

## Introduction

Antiretroviral therapy prevents HIV-1 viral replication but does not impact latent proviruses that are integrated into the genome of host CD4^+^ T cells. The reservoir of latent proviruses is responsible for rapid rebound viremia in most individuals undergoing treatment interruption and is the key impediment to HIV-1 cure ^1–9^.

Although the precise composition of the latent compartment is not known, the relative representation of expanded clones of CD4^+^ T cells harboring intact and defective latent proviruses increases over time such that they account for at least 50 % of the reservoir in chronically infected individuals ^10–17^. Members of infected clones express the same unique T cell receptor (TCR) and have a single distinctive proviral integration site each of which can serve as a molecular identifier for the latent clone ^15, 18–21^. Hypotheses about how the latent reservoir is maintained include proviral integration sites that enable cell division ^22, 23^, homeostatic ^24^ and antigen driven proliferation ^20, 25–28^.

Because cells harboring intact latent proviruses are rare ^5–7, 29, 30^, and have no well-defined markers that distinguish them from other CD4^+^ T cells ^31, 32^, characterizing their transcriptional program has not been possible to date. Intact proviruses are enriched among CD4^+^CD45RA^-^ HLA^-^DR^+^ memory T cells ^24, 33–37^, but other surface markers, such as CD2 ^38^ remain controversial. Latent cells can be identified after re-activation of HIV-1 transcription *in vitro*^18, 39–43^, and there are numerous cell line or tissue culture-based models of latency ^38, 44–48^, but how well these experimental conditions and latency models reflect the physiology of latent cells in circulation is not known.

Here we present a strategy to enrich clones of quiescent latent cells from samples that were assayed directly *ex vivo* from 6 individuals living with HIV-1 based on cell surface expression of their unique TCRs. The enrichment strategy enabled analysis of the transcriptional landscape of these rare cells and identification of distinct features of this population.

## Results

During development T and B lymphocytes assemble unique cell surface receptors by variable diversity and joining gene (V(D)J) recombination. This process is under feedback regulation by the receptor such that each lymphocyte expresses a single specificity, a phenomenon referred to as allelic exclusion ^49^. Because antigen receptors are fixed early in development, naïve T cells that become activated and expand produce clones of CD4^+^ T cells that are defined by expression of a singular TCR. Thus, the TCR expressed by a clone of latent cells is a unique molecular identifier for members of that clone, and because it is a cell surface protein, the TCR can be used to enrich members of the latent clone.

People living with HIV-1 frequently harbor large, expanded clones of latent CD4^+^ T cells. We studied 6 chronically infected individuals controlled on anti-retroviral therapy that were aviremic at the time of sample collection (Table S1). Near-full length sequencing, envelope gene (*env)* sequencing, and/or viral outgrowth assays showed that the latent reservoir of each of these individuals was dominated by a single expanded intact latent proviral clone (Table S2)^11, 18, 21, 26, 50–52^. In all cases the members of these clones could be identified by the sequence of their *env* gene (Figure S1). The frequency of the clonally expanded latent provirus of interest in the 6 individuals ranged from 13 - 431/10^6^ total CD4^+^ T cells based on the frequency of the specific HIV-1 *env* sequence (Table S2).

Latent cells are predominantly found in the CD45RA^-^ memory T cell compartment ^24, 53^. To determine whether the CD4^+^ T cells harboring latent proviral clones of interest are in the memory compartment we purified CD45RA^+^ and CD45RA^-^ cells and analysed proviral DNA by sequencing the *env* gene and producing maximum likelihood phylogenetic trees. As expected, in all cases the latent clone of interest was predominantly found in the CD45RA^-^ population (Figure 1A and Figure S2A). Based on staining with an anti-CD45RA antibody and flowcytometric analysis of CD4^+^ T cells from five individuals, CD45RA^-^ cells accounted for 42 - 68 % of all CD4^+^ T cells and therefore purification of CD45RA^-^ cells results in a 1.5 - 2.4-fold enrichment of the latent clone (Figure S2B).

**Figure 1.**
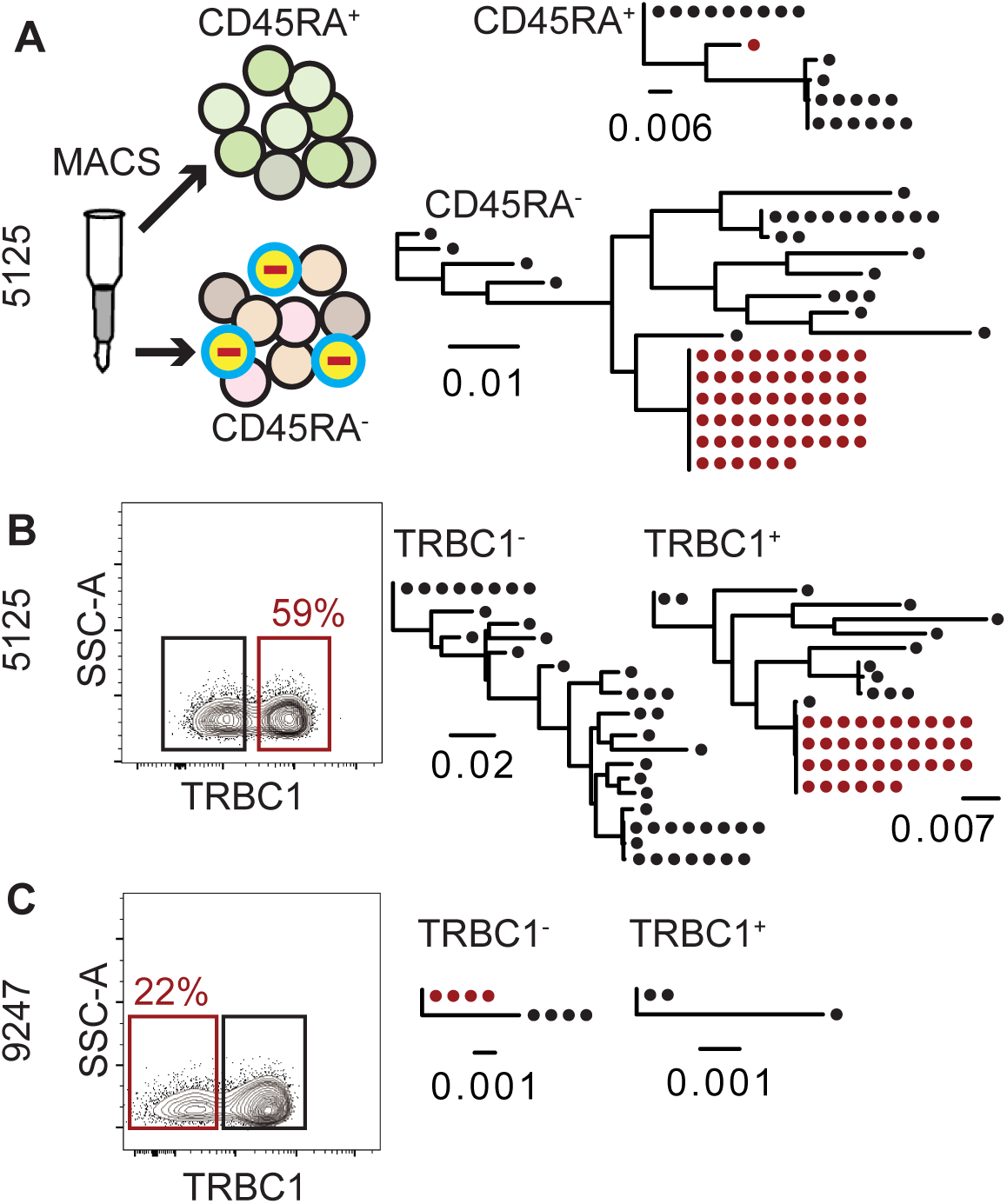
Screening for intact latently infected cell enrichment by sorting for CD45RA and TRBC. Red outlines indicate the population containing the clone of interest. Maximum-likelihood phylogenetic trees show *env* gene of the latent clone of interest marked in red. Each dot represents a recovered *env* sequence from the respective subpopulation of CD4^+^ T cells. The scale bars indicate the number of substitutions per site. (A) CD4^+^ T cells from individual **5125** were magnetically sorted into a CD45RA^+^ and CD45RA^-^ population. Subsequent gDNA extraction, limiting dilution, and *env* sequencing revealed that the infected clone of interest in was enriched in the CD45RA^-^ memory compartment. (B) CD4^+^ T cells from individual **5125** were stained with anti-TRBC1 antibody and sorted into TRBC1^+^ and TRBC1^-^ populations. The latent clone was enriched in the TRBC1^+^ compartment. (C) CD4^+^ T cells from individual **9247** were stained with anti-TRBC1 antibody and sorted into TRBC1^+^ and TRBC1^-^ populations. The latent clone was enriched in the TRBC1^-^ compartment.

The TCRβ locus encodes 2 different constant region genes (TRBC1 and TRBC2). Allelic exclusion ensures that all members of a CD4^+^ T cell clone express the same TCRβ constant region. To determine which of the 2 different TCRβ constant regions is expressed by each of the latent clones of interest we performed flow cytometry experiments to purify TRBC1^+^ and TRBC1^-^ CD4^+^ T cells. In each case, *env* sequencing revealed that the latent provirus was found among CD4^+^ T cells expressing one of the two TRBC domains resulting in 1.5 - 4.5-fold enrichment (Figure 1B and C, and Figure S3).

The TCRβ locus contains 48 functional variable domains (TRBV). To determine whether the TRBV can be used to enrich quiescent clones of latent cells, we made use of a collection of 24 different anti-TRBV monoclonal antibodies. The antibodies were divided into 8 groups of 3 each that were conjugated with either phycoerythrin (PE), or fluorescein isothiocyanate (FITC), or both (PEFITC).

The TRBV expressed by CD4^+^ T cells harboring the latent HIV-1 proviral clones of interest in individuals 603, 605, and B207 were known (TRBV-19, TRBV11-2 and TRBV7-8 respectively)^18^. However, the TRBV expressed by CD4^+^ T cells harboring the latent HIV-1 proviral clones of interest in individuals 5104, 5125, and 9247 were not known. To identify the TRBV expressed by CD4^+^ T cells that harbor the latent clone in these individuals we combined limiting dilution cell sorting with *env* sequencing. As a first screening step, CD4^+^ T cells were stained with the 24 anti-TRBV antibodies and sorted into TRBV^+^ and TRBV^-^ populations. The latent clone in individual 5104 was present in the TRBV^-^ population, yielding a 2-fold enrichment (Figure 2A). For individuals 5125 and 9247, the latent clone was found in the TRBV^+^ population (Figure 2B and C). To identify the precise TRBV expressed by the latent clone in individual 9247, the 24 anti-TRBV antibodies were split into 2 groups of 12 antibodies and CD4^+^ T cells were stained with either one of the 2 groups (Figure 2D). The latent clone was found among one of the 2 groups and the 12 antibodies split again into 4 groups of 3 anti-TRBV antibodies, each group containing one FITC-, one PE-, and one PEFITC-labelled antibody (Figure 2E). The latent clone was only found in one of the 4 groups, and the 3 anti-TRBV antibodies in that group were then used to stain and sort for each of the anti-TRBV individually (Figure 2F). These experiments revealed that in individual 9247, the latent clone expresses TRBV4-3 (Figure 2F). For individual 5125, the 24 TRBV antibodies were split into 8 groups of 3 and the latent clone was found in only one of the groups (Figure 2G). The 3 antibodies in that group were then used to purify the individual TRBV expressing cells, which showed that the latent clone of interest was found among TRBV2^+^ cells (Figure 2H).

**Figure 2.**
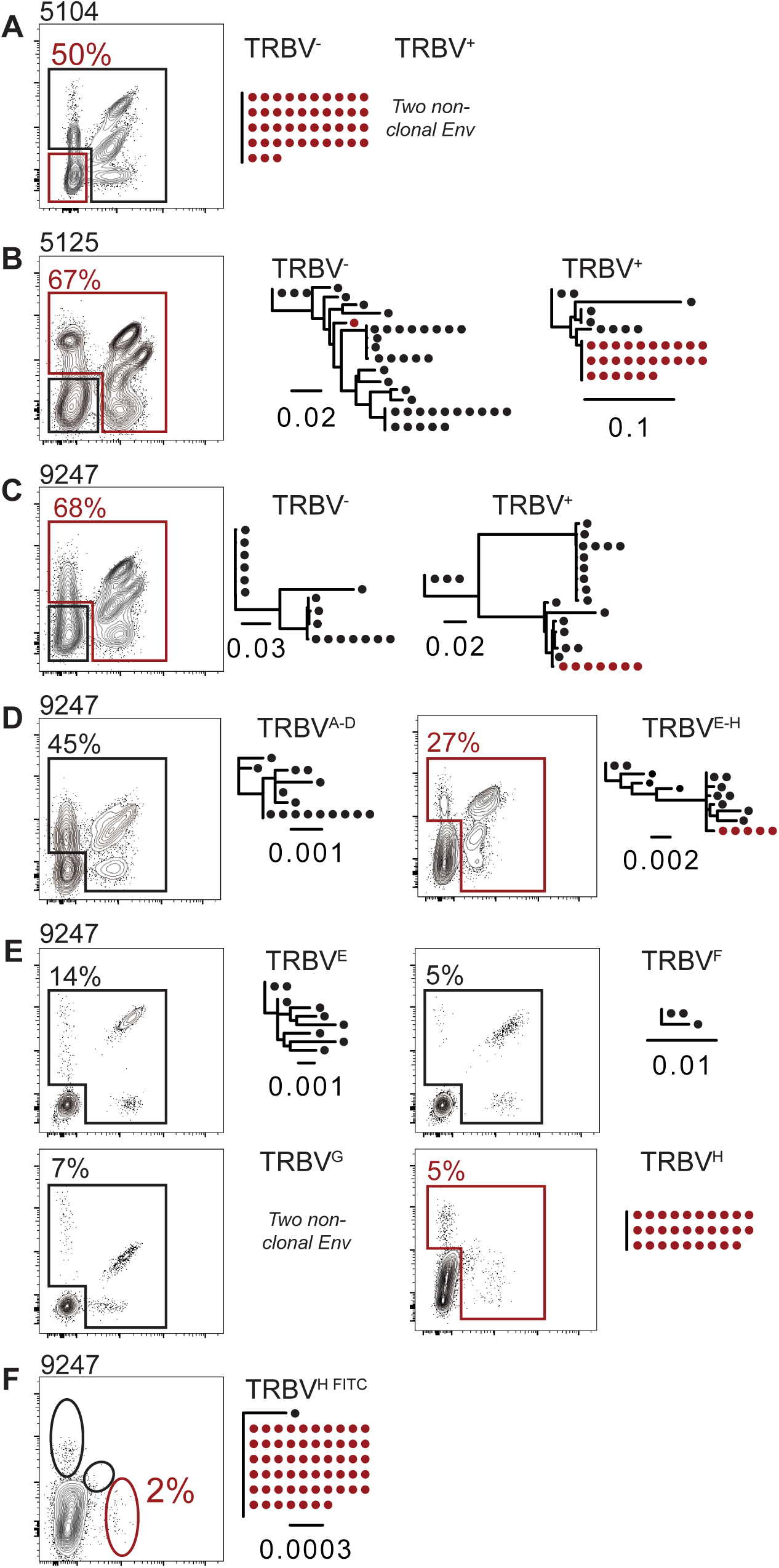
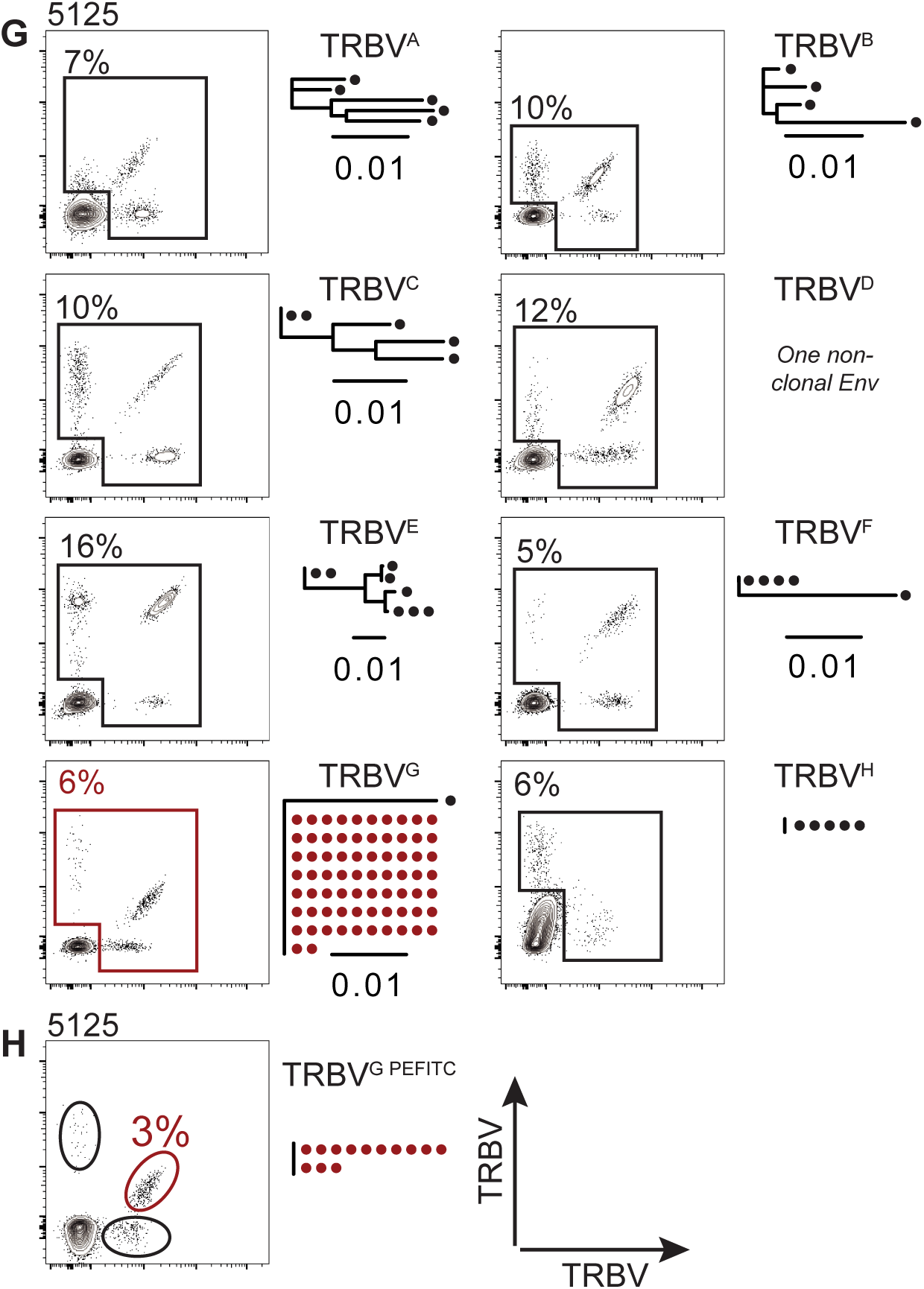
Screening for the TRBV expressed by the latent clone in individuals 5104, 5125 and 9247. Red outlines indicate the population containing the clone of interest. Maximum-likelihood phylogenetic trees of *env* sequences derived from sorted population, as indicated. *Env* gene of the clone of interest marked in red. The scale bars indicate the number of substitutions per site. TRBV^A-H^ indicate the grouping of monoclonal antibodies used in the cocktails (see methods). (A-C) Flow cytometry plots show TRBV staining using a combination of 24 anti-TRBV antibodies. Percentage of TRBV^-^ cells is indicated for individual **5104** and TRBV^+^ cells for **9247** and **5125**. (D) Flow cytometry plots show TRBV staining with one of 2 groups of 12 antibodies for individual **9247**. Percentages of TRBV^+^ cells are indicated. The sorted population is indicated. The positive group was further subdivided in (E). (E) Flow cytometry plots show TRBV staining with one of 4 groups of 3 TRBV antibodies from individual **9247**. Percentages indicate the fraction of TRBV^+^ cells in each group. The sorted population is indicated. The clone of interest was only found in cells stained with the TRBV^H^ cocktail indicated in red and was further subdivided in (F). (F) Flow cytometry plots show TRBV staining with 3 different monoclonal antibodies from individual **9247**. Percentage indicates the fraction of TRBV4-3^+^ cells. (G) Flow cytometry plots show TRBV staining with antibody cocktails A-H from individual **5125**. The clone of interest was only found in the TRBV^G^ cocktail which was further subdivided in (H). Percentage indicates the fraction of TRBV^+^ cells for each cocktail. (H) Flow cytometry plots show TRBV staining with the 3 different antibodies in the TRBV^G^ cocktail from individual **5125**. The clone of interest was found in the TRBV2^+^ population. Percentage indicates the fraction of TRBV2^+^ cells.

To determine the amount of enrichment that could be achieved for each individual we combined anti-CD45RA, -TRBC, and -TRBV enrichment and performed limiting dilution, *env* amplification and sequencing on genomic DNA (Figure 3 and Table S3).

**Figure 3.**
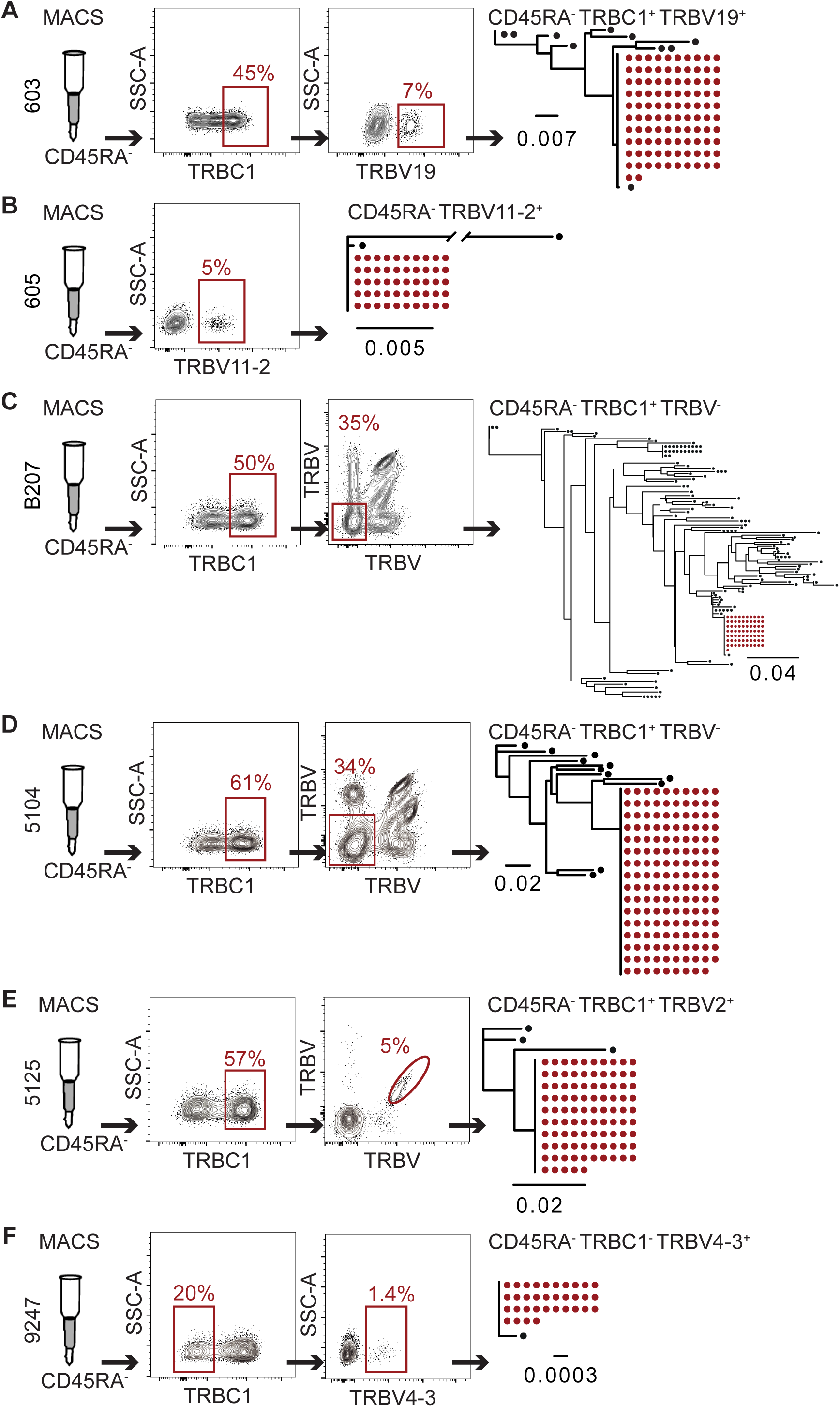
Combined enrichment based on CD45RA, TRBC, and TRBV expression in individuals 5104, 5125, and 9247. Red outlines indicate the population containing the clone of interest. Maximum-likelihood phylogenetic trees of *env* sequences derived from sorted population, as indicated. *Env* gene of the clone of interest marked in red. The scale bars indicate the number of substitutions per site. (A) Enrichment in individual **603** after purification of CD45RA^-^ memory cells followed by staining and sorting for TRBC1^+^ and TRBV19^+^ cells. Flow cytometry plots show TRBC1 and TRBV19 staining. Percentages of CD45RA^-^TRBC1^+^, and CD45RA^-^TRBC1^+^TRBV19^+^ cells are indicated. (B) Enrichment in individual **605** after purification CD45RA^-^ memory cells followed by staining TRBV11-2^+^ cells, which were > 90 % TRBC1^+^ (Figure S3B). Flow cytometry plot shows TRBV11-2 staining. Percentage of CD45RA^-^TRBV11-2^+^ cells is indicated. (C) Enrichment in individual **B207** after purification of CD45RA^-^ memory cells followed by staining and sorting for TRBC1^+^ and TRBV^-^ cells. Flow cytometry plots show TRBC1 and combined 24 TRBV staining. Percentages of CD45RA^-^TRBC1^+^, and CD45RA^-^TRBC1^+^TRBV^-^ cells are indicated. (D) Enrichment in individual **5104** after purification CD45RA^-^ memory cells followed by staining and sorting for TRBC1^+^ and TRBV^-^ cells. Flow cytometry plots show TRBC1 and TRBV staining. Percentages of CD45RA^-^TRBC1^+^, and CD45RA^-^TRBC1^+^TRBV^-^ cells are indicated. (E) Enrichment in individual **5125** after purification CD45RA^-^ memory cells followed by staining and sorting for TRBC1^+^ and TRBV2^+^ cells. Flow cytometry plots show TRBC1 and TRBV2 staining. Percentages of CD45RA^-^TRBC1^+^, and CD45RA^-^TRBC1^+^TRBV2^+^ cells are indicated. (F) Enrichment in individual **9247** after purification of CD45RA^-^ memory cells followed by staining and sorting for TRBC1^+^ and TRBV4-3^+^ cells. Flow cytometry plots show TRBC1 and TRBV4-3 staining. Percentages of CD45RA^-^TRBC1^-^, and CD45RA^-^TRBC1^-^TRBV4-3^+^ cells are indicated.

To enrich the latent clone in individual 603, CD4^+^ T cells were magnetically sorted for CD45RA^-^cells and then stained with antibodies to TRBC1 and TRBV19. The latent clone was found in the CD45RA^-^TRBC1^+^TRBV19^+^ population resulting in an overall 47-fold relative enrichment (Figure 3A and Table S3). The latent clone in individual 605 was enriched by 40-fold in the CD45RA^-^ TRBV11-2^+^ population (Figure 3B, Table S3, and Figure S3B). Antibodies to TRBV7-8 expressed by the latent clone in individual B207 were not available. Therefore, B207 CD4^+^ T cells were stained with the 24 anti-TRBV antibodies and sorted into a CD45RA^-^TRBC1^+^TRBV^-^ population resulting in only a 9-fold overall enrichment (Figure 3C and Table S3). The latent clone in individual 5104 was found in the CD45RA^-^TRBC1^+^TRBV^-^ population with an 11-fold enrichment (Figure 3D and Table S3). The latent clone in individual 5125 was found in the CD45RA^-^TRBC1^+^TRBV2^+^ population with a 54-fold enrichment (Figure 3E and Table S3). Lastly, the latent clone in individual 9247 was found in the CD45RA^-^TRBC1^-^ TRBV4-3^+^ population with a 675-fold enrichment (Figure 3F and Table S3).

After combined enrichment the latent provirus of interest is found in 20, 163, 6, 14, 15 and 18 in 10^4^ CD4^+^ T cells in individuals 603, 605, B207, 5104, 5125, and 9247 respectively (Table S4).

Although we were able to identify the TRBV expressed by expanded clones of interest in individuals 5104, and 5125, and 9247, the precise TCRαβ sequence remained unknown. Candidate TCRs were initially identified among clones of CD4^+^ T cells in the enriched populations by single cell TCR sequencing by 10x Genomics (Figure 4A). To definitively determine the TCR expressed by the clone of CD4^+^ T cells harboring the latent provirus we combined anti-CD45RA, -TRBC, and -TRBV staining and sorted 5 candidate cells into multi-well plates, followed by DNA and RNA extraction. *Env* amplification and sequencing from genomic DNA was used to identify wells containing latent cells. cDNA from wells containing the latent provirus or negative controls was used to amplify and sequence TCRα and β chains (Figure 4A). CD4^+^ T cells from individuals 603 and 605 that express known TCRs were used as positive controls to validate the method (Figure 4B and C). For individual 603, the TCR expressed by the latent clone was found in 63 % of Env^+^ wells and only in 4 % of Env^-^ wells (Figure 4B). In 605, the specific TCR was found in 90 % of the Env^+^ wells and only in 6 % of Env^-^ wells. In both cases there was no enrichment of irrelevant TCRs in Env^+^ wells. All 7 Env^+^ wells obtained from individual 5104 contained TCRα TRAV12-3/J9 and/or TCRβ TRBV5-4/J1- 1 representing a unique TCR clone in the 10x Genomics sequencing data (Figure 4B and C).

**Figure 4.**
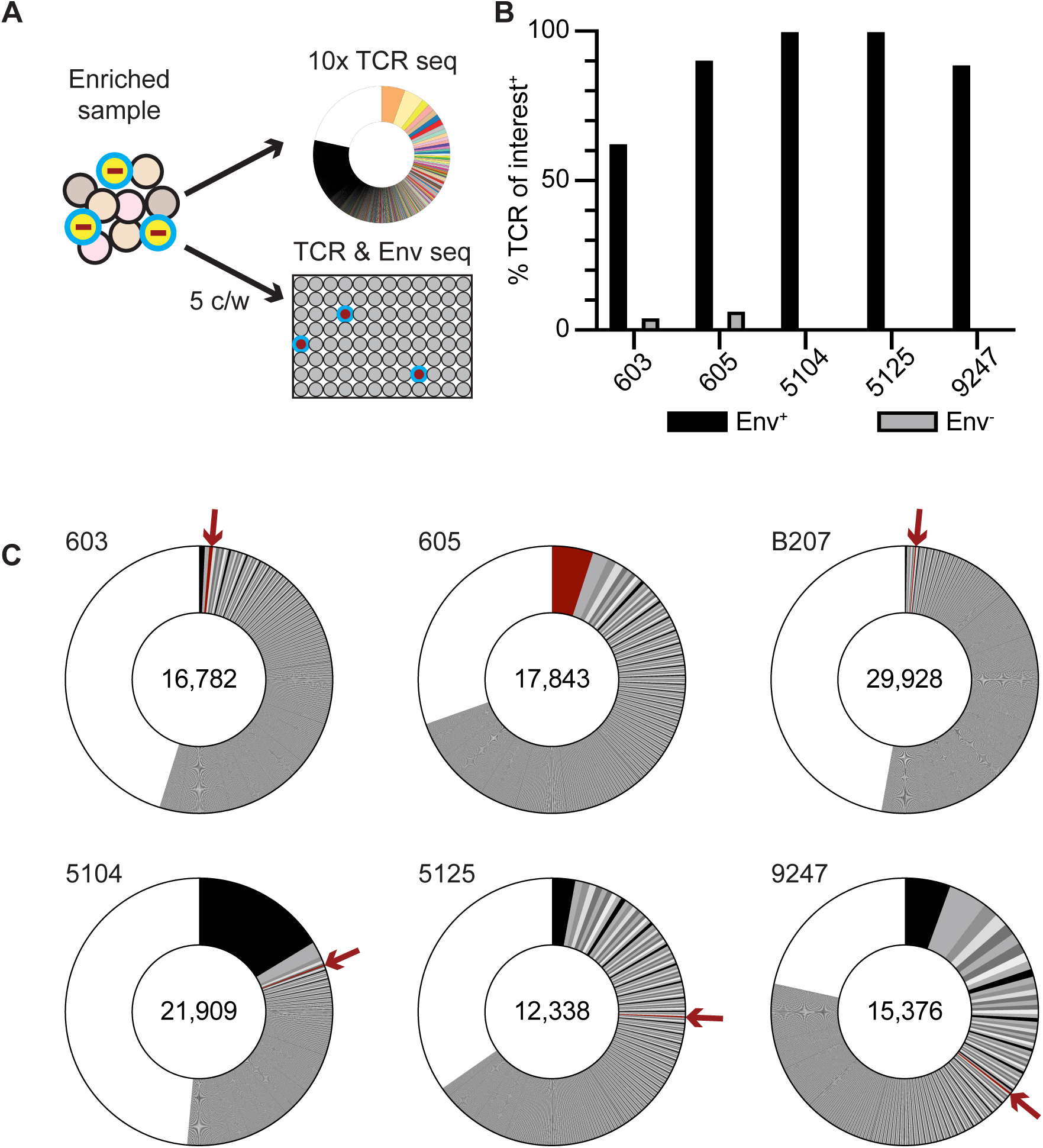
Identification of latent clone TCR sequences. (A) Limiting dilution sorting strategy to identify the TCR expressed by the CD4^+^ T cells harboring the clone of interest. Each sample was enriched based on CD45RA, TRBC1, TRBV expression, and 5 cells sorted per well (c/w) into microwell plates. *Env* sequencing identified wells that contained a cell of the latent clone of interest. TCRs were sequenced from all *env*^+^ and a selection of *env*^-^ wells. (B) Bar graph shows the relevant TCR clonotypes identified in *env*^+^ and *env*^-^ wells. **603**: *env*^+^ n=8 and *env*^-^ n=77; **605**: *env*^+^ n= 21 and *env*^-^ n=62; **5104**: *env*^+^ n=7 and *env*^-^ n=36; **5125**: *env*^+^ n=4 and *env*^-^ n=31; **9247**: *env*^+^ n=9 and *env*^-^ n=42. (C) Pie charts show the relative size of TCR clones as slices. The areas indicated in white represent unique TCR sequences. The number on the left above the pie chart is the donor ID for each individual. The number in the center of the pie chart represents the number of cells assayed for each individual. The clone of interest is indicated by a red arrow and pie slice.

This TCR was absent in the random selection of Env^-^ wells. Similarly, all 4 Env^+^ wells from individual 5125 contained TRAV26-2/J32 and/or TRBV2/J1-1, that was not found among Env^-^ wells. Finally, in individual 9247 TRAV38-1/J33 and/or TRBV4-3/J2-3, was present in 8 out of 9 of Env^+^ wells but absent in the random selection of Env^-^ wells. We conclude that the latent clone of interest in 5104, 5125 and 9247 express TRAV12-3/J9/TRBV5-4/J1-1, TRAV26- 2/J32/TRBV2/J1-1, and TRAV38-1/J33/TRBV4-3/J2-3 respectively (Figure 4C).

HIV-1 proviruses can integrate into CD4^+^ T cells undergoing clonal expansion at the time they start dividing or sometime thereafter. Proviral integration in early stages of clonal expansion would yield a homogenous group of cells the vast majority of which would harbor an HIV-1 provirus in the same genomic location. Integration at a later time would produce a heterogeneous CD4^+^ T cell clone wherein only some of the cells in the expanded clone harbor the HIV-1 provirus ^20^. To estimate the fraction of infected cells within a particular clone based on its representation by TCR, we performed 10x Genomics single cell TCR sequencing on samples enriched using the antibody methods described above. We compared the TCR frequencies to the relative frequency of the specific *env* from proviral DNA in similarly enriched samples (Table S4). For individuals 5104, 5125, and 9247, the frequency of the specific provirus was similar to the frequency of corresponding TCR, but in B207, 603, and 605, the number of cells expressing the specific TCR of interest was 2-3 times higher than the frequency of proviral copies (Table S4). Thus, there is heterogeneity among clones of expanded CD4^+^ T cells that harbor latent HIV- 1 proviruses. In half of our samples most clonally expanded CD4^+^ T cells harbor latent HIV-1, and in the others the provirus is found in a fraction of the clone.

To determine whether CD4^+^ T cell clones harboring latent proviruses share a transcriptional profile we combined the 10x Genomics mRNA and TCR sequencing data obtained from the enriched populations of latent cells. TCR sequencing data was used to identify CD4^+^ T cell clones harboring the latent HIV-1 provirus but was omitted for gene expression analysis to prevent TCR-biased clustering. Uniform manifold approximation and projection (UMAP) analysis of the transcriptional profile of all the 109,217 cells from the six individuals produced 15 unique clusters of CD4^+^ memory T cells (Figure 5A). Visual inspection revealed that CD4^+^ T cells expressing the specific TCR associated with latent proviruses are found predominantly in gene expression cluster 7 and neighbouring clusters (Figure 5A). On average 57 % of all cells expressing the TCR associated with the expanded latent clone in the 6 individuals were found in cluster 7. Except for individual 9247, the fraction of CD4^+^ T cells belonging to the latent clone in cluster 7 was greater than all other clusters ranging from 48 - 73 % (Figure 5B and Table S5). In individual 9247, the largest fraction of latent cells was in cluster 6 (35 %), which is closely related to cluster 7, and the second largest fraction of latent cells was in cluster 7 (29 %).

**Figure 5.**
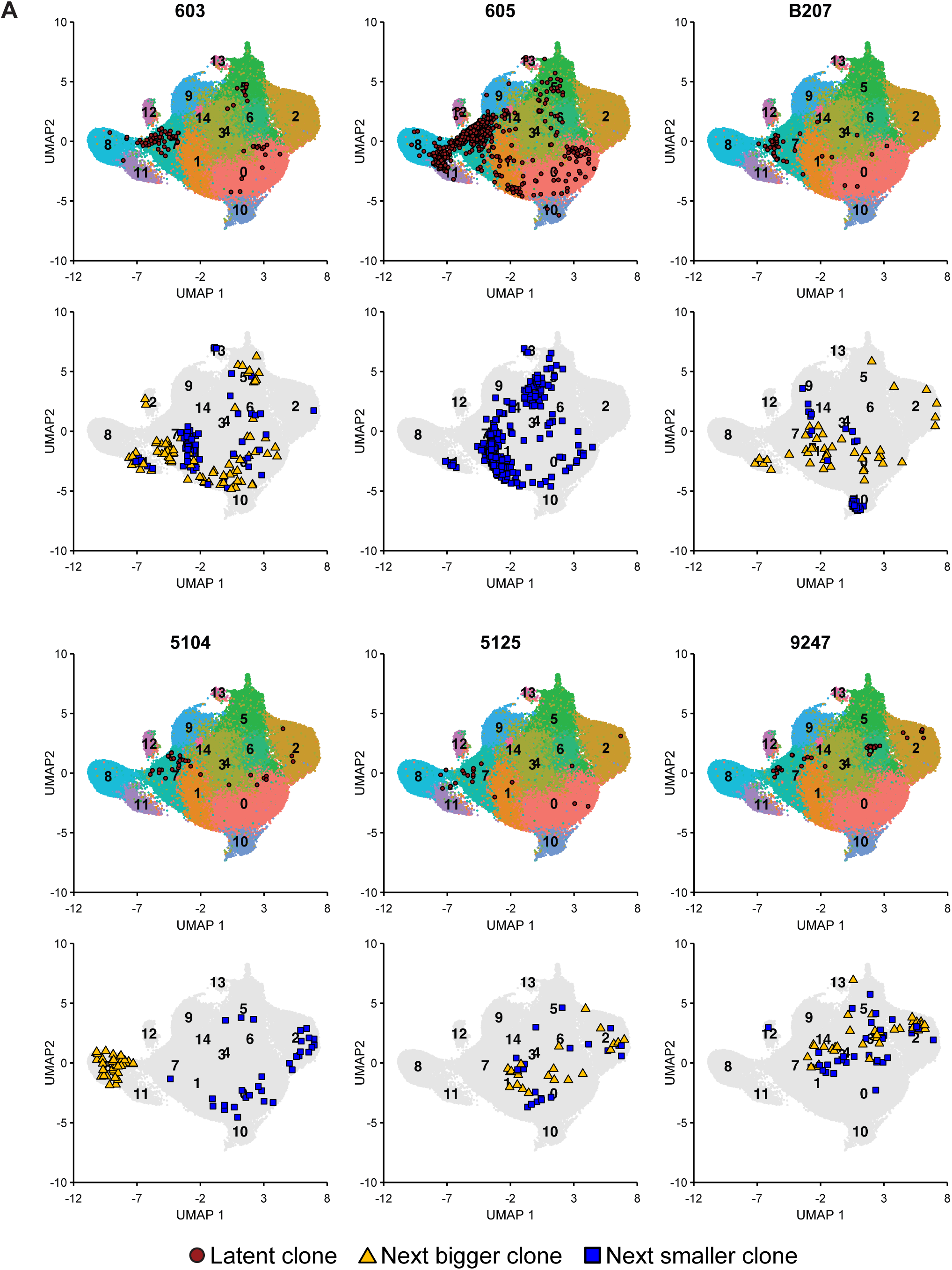
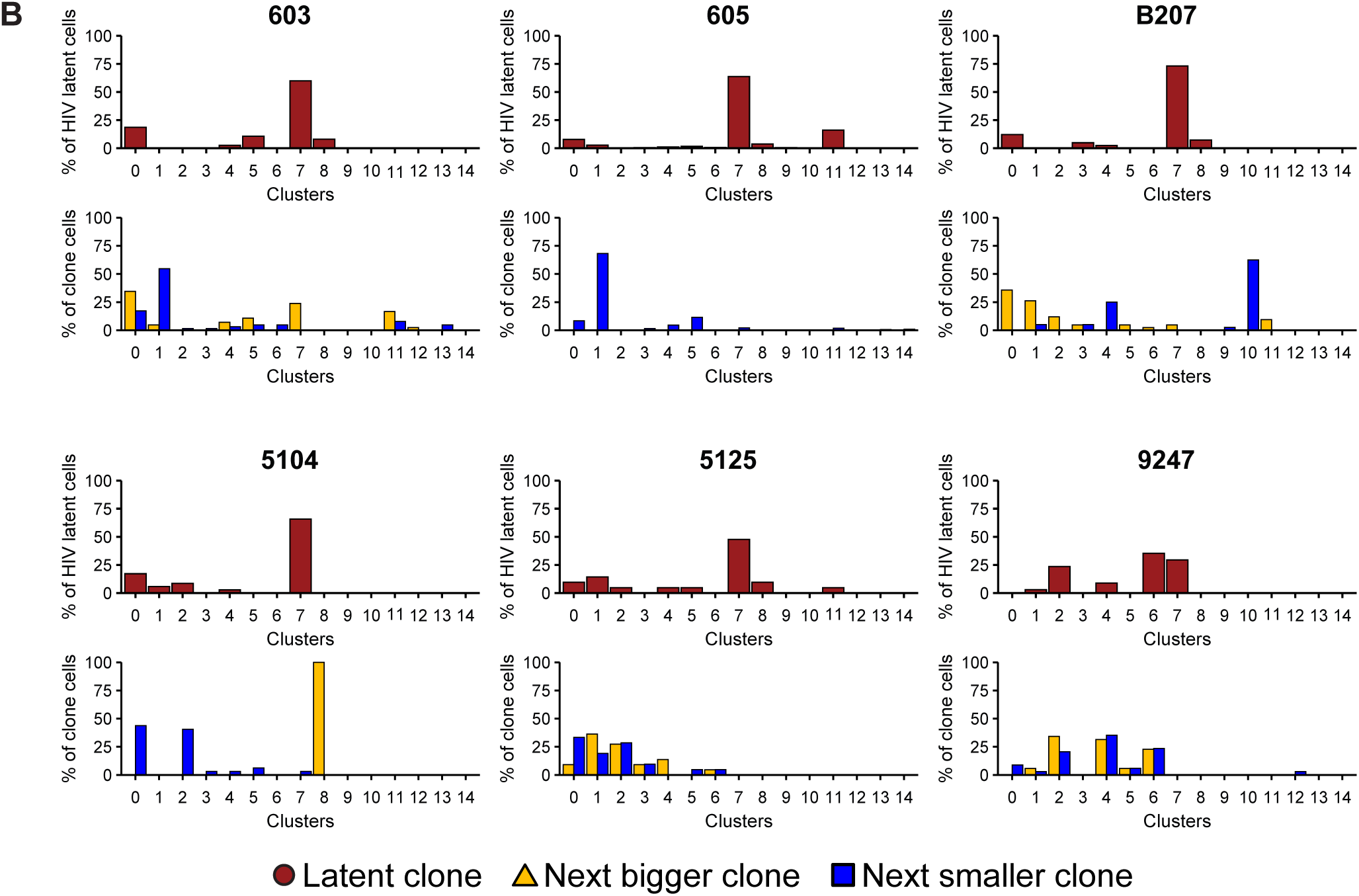
Uniform Manifold Approximation and Projection (UMAP) of 10x gene expression data. Data representing mRNA expression by 109,217 individual cells is shown. The latent clone of interest as well as the next bigger and next smaller clone in size were located in the UMAP by their TCR sequence. (A) UMAPs show the position of the cells expressing the latent clone TCR for each of the 6 individuals as red dots. Underneath, UMAPs show the position of the next bigger (yellow triangles) and the next smaller (blue squares) clone in size to the clone of interest. For individual **605**, the latent clone of interest was the biggest clone, and only the next smaller clone is shown. (B) The bar graphs show the fraction of cells in the latent clone (red bars) in each of the 15 UMAP clusters, and the fraction of cells of the next bigger (yellow bars) and the next smaller (blue bars) clone in size to the clone of interest in each of the 15 UMAP clusters (Table S3).

To determine whether residence in cluster 7 is a general property of expanded clones of memory CD4^+^ T cells, we determined the position of the cells in the next largest and/or smallest clones of CD4^+^ memory T cells in the UMAP (Figure 5A). None of the 11 neighbouring CD4^+^ T cell clones examined were predominantly found in gene expression cluster 7. In addition, some TCR clones clustered by gene expression whereas others did not. For example, in individual B207, cells in the next largest clone to the one containing latent proviruses were found in 8 of the 15 clusters. In contrast, cells in the next smaller clone in B207 were found primarily in gene expression cluster 10 (63 %) which contains cells expressing Foxp3 (Figure 5B, Table S5). We conclude that clonal expansion per se is not sufficient for CD4^+^ memory T cell accumulation in gene expression cluster 7.

Examination of the top 120 genes that define the UMAP clusters revealed that cluster 7 and 8 are closely related (Figure 6A). Cluster 7 is enriched in genes that encode antigen-presenting molecules or their chaperones such as HLA-DR and HLA-DP, as well as CD74, the invariant chain for major histocompatibility class II (MHCII) molecules ^54^ (Figure 6A and Table S6). In addition, cluster 7 is distinguished by expression of CCL5, Granzymes A and K (GZMA, GZMK), cystatin F (CST7), and the nuclear proteins LYAR ^55, 56^ and DUSP2 ^57, 58^. Differential expression of some of these genes has been reported in activated latent CD4^+^ T cells ^18, 36, 37^. However, Granzyme A, and K proteins do not accumulate specifically in latent cells because flow cytometry-based cell sorting for these markers did not enrich latent cells.

**Figure 6.**
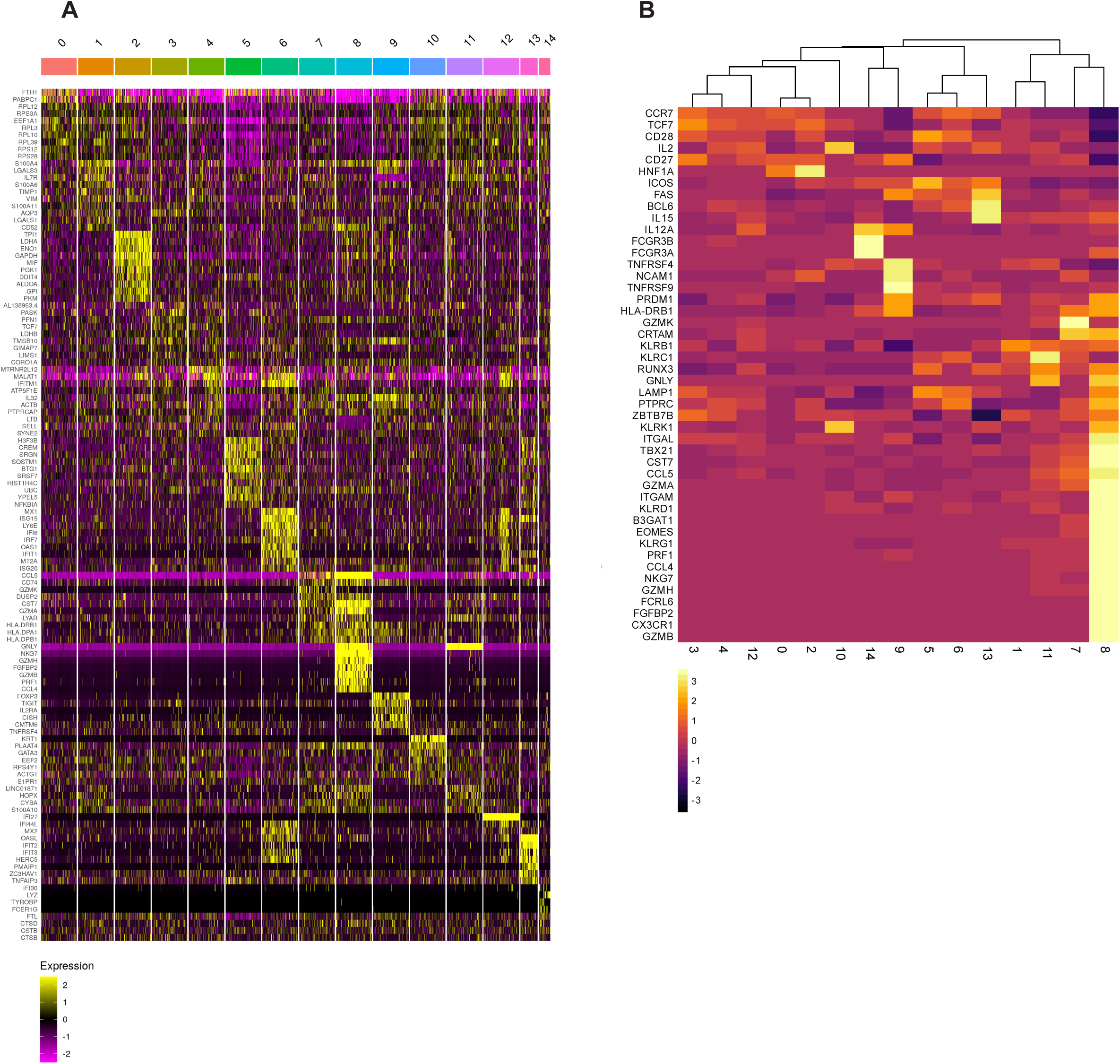
Heat map of cluster-defining, differentially expressed genes. (A) Heat map shows up to 10 of the most up-regulated genes per UMAP cluster compared to all other clusters. Genes are indicated on the left and clusters above. Yellow shows relatively highly expressed genes and purple relatively downregulated genes. (B) Heat map shows unsupervised clustering based on the expression of differentially expressed genes in cytotoxic CD4^+^ T cells ^60, 61, 63–66^. Genes are indicated on the left and clusters below. Yellow shows relatively highly expressed genes and black shows relatively downregulated genes.

Cluster 7 shares many upregulated genes with cluster 8, namely CCL5, CST7, and GZMA (Figure 6A). The two closely related clusters differ in that cluster 8 cells also express GZMB, GZMH, perforin 1 (PFR1), natural killer cell granule protein 7 (NKG7), and granulysin (GNLY), each of which is associated with cytotoxic CD4^+^ T cells ^59, 60^,. This population of cells is frequently expanded in infection and chronic inflammation ^61^ and is also found enriched among tumour infiltrating lymphocytes ^62^. Moreover, the relative proportion of cytotoxic CD4^+^ T cells among memory cells is expanded in HIV-1 infected individuals including those on suppressive ART^63, 64^.

To examine the relationship between the cells in UMAP clusters and genes associated with CD4^+^ T cell identity we performed unsupervised clustering analysis with a collection of genes that are up or downregulated in these cells ^60, 61, 63–66^ (Figure 6B). In agreement with the UMAP, cluster 8 stands out as most closely related to CD4^+^ cytotoxic cells and cluster 7 is its closest relative. Nonetheless, cluster 7 differs from cytotoxic CD4^+^ T cells in cluster 8 in several respects including expression of higher levels of CD27, CD28, and CCR7 ^63^, and lower levels of the transcription factors eomesodermin (EOMES) and RUNX3 ^61, 65, 66^.

To further characterize the expanded clones of latent cells harboring intact HIV-1 proviruses, the 10x gene expression data was projected on a reference data set of a multimodal single cell analysis of PBMCs from HIV-1 negative individuals ^67^. Cluster 8 falls into the cytotoxic CD4^+^ T cell population, and cluster 7 falls into the CD4^+^ T effector memory population (T_EM_) that is marked by the expression of granzymes A and K, DUSP2, CST7, LYAR, and HLA-DRB1 (Figure 7) ^67^. In four individuals, the overall fraction of cells in the clones harboring intact HIV-1 proviruses was greatest in the T_EM_ compartment (Figure 7). In two other individuals, 5104 and 9247, the clones were enriched in the central memory compartment (T_CM_). However, the T_CM_ population is larger than the T_EM_ compartment, and when corrected for compartment size, all individuals but individual 605 showed relative enrichment of the latent clone in among T_EM_ cells (Table S7).

**Figure 7.**
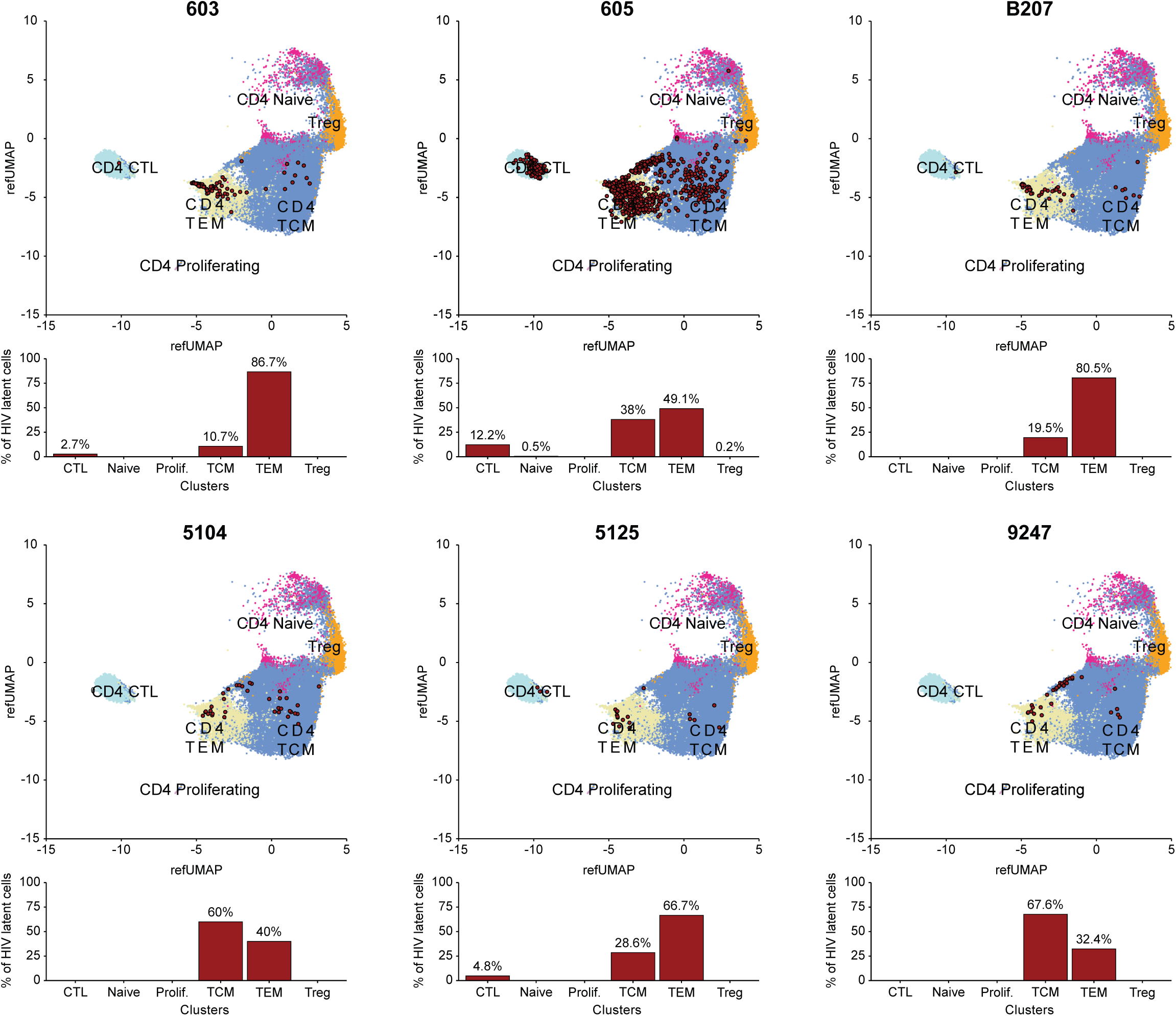
Projection of 10x gene expression data on Uniform Manifold Approximation and Projection (UMAP) of CD4^+^ T cells from multimodal single cell sequencing. ^67^ Projection of data representing mRNA expression by 109,217 individual cells on a multimodal UMAP of CD4^+^ T cells from HIV-negative individuals. The latent clone of interest in each individual was located in the UMAP by its TCR sequence and is represented as red dots. Underneath each UMAP, the bar graph shows the fraction of cells in the latent clone in each T cell subpopulation as indicated by the number above each bar.

In conclusion, gene expression cluster 7 is characteristic of CD4^+^ T effector memory cells. This cluster harbors some of the features of cytotoxic CD4^+^ T cells but differs from cytotoxic cells in the expression in key transcriptional factors and components of the cytotoxic machinery ^61, 66^.

## Discussion

CD4^+^ T cells harboring intact latent HIV-1 proviruses represent only a tiny fraction of all CD4^+^ T cells. They are usually found among subsets of the memory cells ^24, 53^ but there are no specific markers that facilitate the purification of these cells ^31, 32^. Consequently, their transcriptional program has only been studied after activation of HIV-1 expression which thereby allows their identification ^18, 39–42^. We have devised a novel method to enrich quiescent latent memory CD4^+^ T cells by means of their specific antigen receptors. We find that expanded clones of memory CD4^+^ T cells that carry intact integrated HIV-1 proviruses are enriched among cells that express a transcriptional program that is found in CD4^+^ T_EM_ cells. Our data is in accordance with studies showing enrichment of genetically intact proviruses in the CD4^+^ T_EM_ compartment but extends previous observations by revealing the transcriptional program of resting latent cells ^33, 68^.

T_CM_ cells are antigen experienced T cells that are CD45RA^-^CD45RO^+^CD27^+^CCR7^+^CD62L^+^ and circulate through secondary lymphoid organs ^24, 69–73^. Upon re-stimulation with cognate antigen T_CM_ secrete interleukin 2 (IL-2) and can differentiate further into T_EM_ that are polarized to secrete specific effector cytokines ^73^. T_EM_ cells are CD45RA^-^CD45RO^+^CD27^-^CCR7^-^ and express chemokine receptors that enable them to home to inflamed tissues ^24, 72, 73^. These cells are more committed to specific T_H_ differentiation lineages than T_CM,_ and secrete effector cytokines or function as CTL upon cognate antigen challenge ^69, 73, 74^. The observation that expanded clones of latent cells are frequently found in the CD4^+^ T_EM_ compartment is consistent with the finding that clones of latent cells and T_EM_ develop in response to chronic viral infection ^20, 26, 72, 74–77^.

CD4^+^ cytotoxic T cells are a subset of T_EM_ whose effector function is killing target cells. Like other T_EM_ they develop in response to chronic antigenic stimulation. They are prominently expanded in HIV-1, CMV, EBV infection, chronic inflammatory diseases and in virally induced malignancies ^61–64, 78^. Their polyfunctional phenotype is most closely associated with the Th1 phenotype, but they can also develop from other T cell lineages. The mechanism that regulates their development is not entirely defined but is associated with expression of T-bet, EOMES, Runx3, and Blimp1, and downregulation of ThPOK, Bcl6 and TCF1 ^61, 65, 66^. In keeping with the finding that clones of CD4^+^ T cells harboring intact latent proviruses cluster in close proximity to CD4^+^ cytotoxic cells, this subset of T_EM_ can respond to antigens found in chronic viral infections such as HIV-1, CMV, and EBV ^20, 26, 27, 47^.

CD4^+^ T cell containing integrated proviruses in individuals on suppressive ART can express HIV-1 RNA, but the majority of these cells harbor defective proviruses ^7, 8, 10, 15, 17, 29, 33, 34, 79–82^. When examined based on HIV-1 RNA expression alone, irrespective of whether the provirus is intact, CD4+ T cells containing integrated proviruses are enriched in Granzyme B expression, suggestive of residence in the CD4 cytotoxic T cell compartment ^83^. Although we find a fraction of latent cells containing intact proviruses in the cytotoxic CD4+ T cell compartment, this is a minority in 5 out of 6 individuals.

Members of a T cell clone expressing the same TCR can adopt different fates depending on several different factors including affinity, antigen concentration and the cytokine milieu ^34, 84, 85^. Consistent with this idea, CD4^+^ T cells expressing the TCR associated with latent proviruses are not entirely limited to a single gene expression cluster. Nevertheless, the observation that a large fraction of the cells in the expanded clones we studied can be found in one specific transcriptional cluster among all memory CD4^+^ T cells stands in contrast to other similar sized expanded clones obtained from the same individuals. The latter are found in several different clusters that diverge between clones and individuals, and many of the cells in these clones are widely dispersed among clusters with different transcriptional signatures. For example, among the 11 random memory CD4^+^ T cell clones of similar size examined in the 6 individuals only 1 in individual 603 showed enrichment above 5 % in cluster 7.

In chronically infected individuals at least 50 % of the cells carrying intact proviruses belong to expanded clones ^11, 12^, each of which can be distinguished by expression of a specific T cell receptor that is associated with a unique proviral integration site ^10, 15, 18–20, 22, 23^. The intact reservoir is dynamic and while the absolute number of cells in the reservoir decreases slowly with a half-life of 4 - 5 years, clonality increases with time after infection in people on suppressive ART ^4, 6, 10, 13, 16, 17, 34^. However, clonal expansion is not a unique feature of CD4^+^ T cells harboring intact proviruses. Defective proviruses are also found predominantly in expanded clones ^10, 17, 86^, and clones of CD4^+^ T cells are prominent among non-HIV-1 infected individuals^87^.

Some of the genes that help define quiescent CD4^+^ T cells in cluster 7 have also been reported to be expressed in latent cells, such as HLA-DR ^18, 34, 36^, or like CD2 or LYAR upregulated in *in vitro* models of HIV infection and latency ^38, 88^. In contrast, CCL5 is downregulated upon latent cell reactivation *in vitro* ^18, 38^, but was upregulated in gene expression cluster 7. Notably, these genes had not been linked to a specific transcriptional program that differentiates latent cells from other CD4^+^ T cells.

Cells expressing the genes that define cluster 7 are not a unique feature of HIV-1 infection and can be found in non-infected individuals ^60, 67^. Therefore, HIV-1 proviral integration and latency per se is not required for T cells to acquire the cluster 7 transcriptional program. Why latent proviruses are enriched in cells expressing this particular program is not known. One possibility among many is that the cluster 7 program favors suppression of HIV-1 gene expression during T cell activation which would permit cell division in the absence of HIV-1 virion production and cell death. An alternative but non-exclusive possibility is that these cells are among the most likely to respond to a chronic infection, and therefore the most likely to be infected and become latent and subsequently undergo clonal expansion in response to a persistent antigen.

Our analysis is limited to large, expanded clones of latent cells in 6 individuals and did not include non-circulating CD4^+^ T cell subsets such as tissue-resident CD4^+^ T cells ^89^. Whether these observations also apply to less expanded or tissue resident populations of latent cells remains to be determined. Moreover, the depth of sequencing available on the 10x platform is also limiting and therefore additional elements of the cluster 7 transcriptional program remain to be defined. Despite these caveats, the observation that latent cells preferentially display a specific transcriptional program suggests that these cells could be specifically targeted for elimination.

---

**Figure S1.**
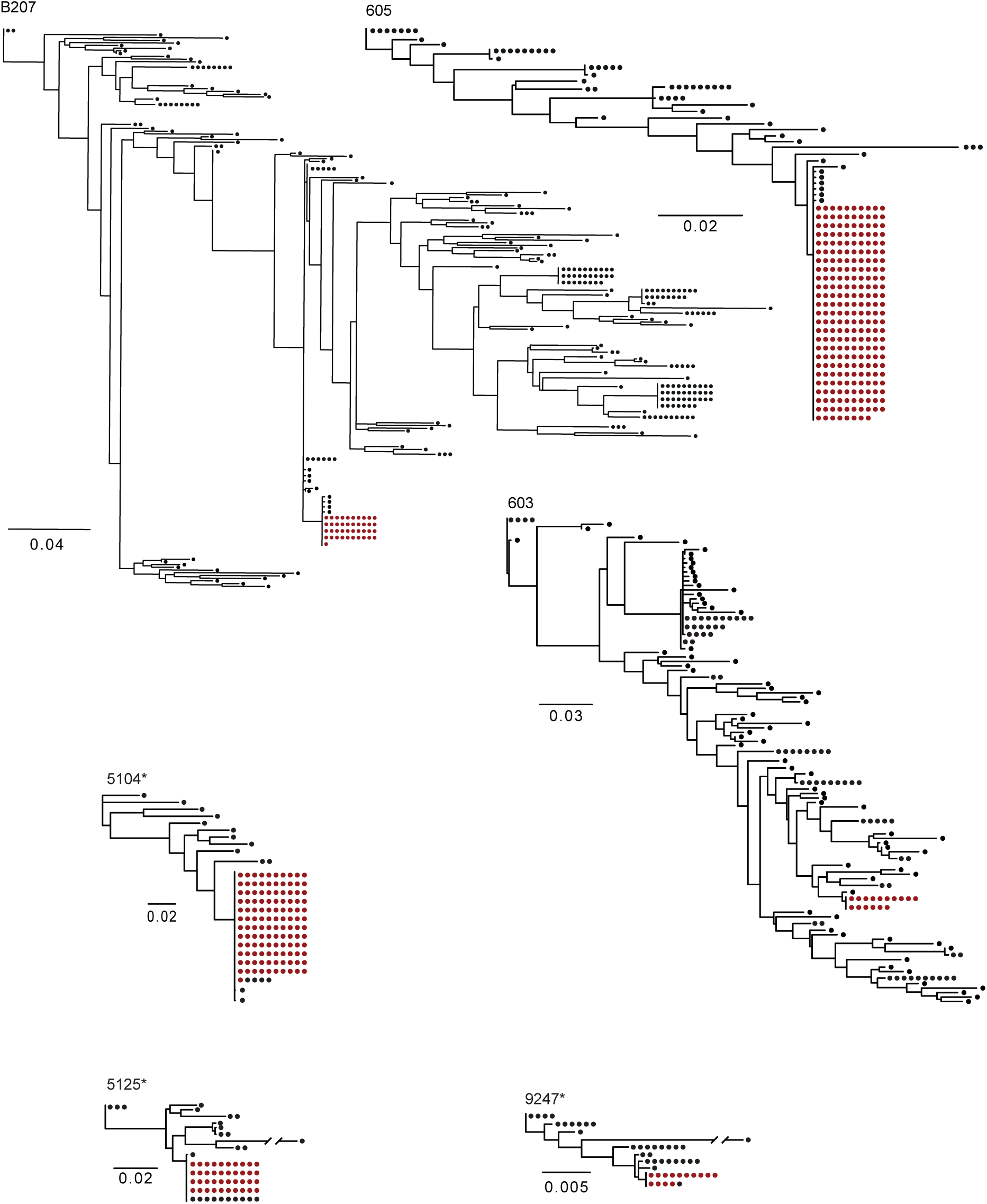
T**h**e **latent reservoir in each individual contains an expanded intact latent clone** Maximum-likelihood phylogenetic trees show *env* gene of the clone of interest marked in red. Each dot on the maximum-likelihood phylogenetic trees represents an *env* sequence that was recovered by *env* PCR (individuals **B207**, **603**, and **605**) or Q4PCR (individuals **5104**, **5125**, and **9247** ^51, 52^) from CD4^+^ T cells before enrichment. The *env* gene that identifies the clone of interest is marked red. The scale bars indicate the number of substitutions per site. For individuals **B207**, **603**, and **605**, the *env* gene was also found in viral outgrowth assays ^18^. For individuals **5104**, **5125**, and **9247**, an expanded intact clone was identified by Q4PCR ^50^. All *env* sequences were extracted from near-full-length proviral sequences and aligned in trees.

**Figure S2.**
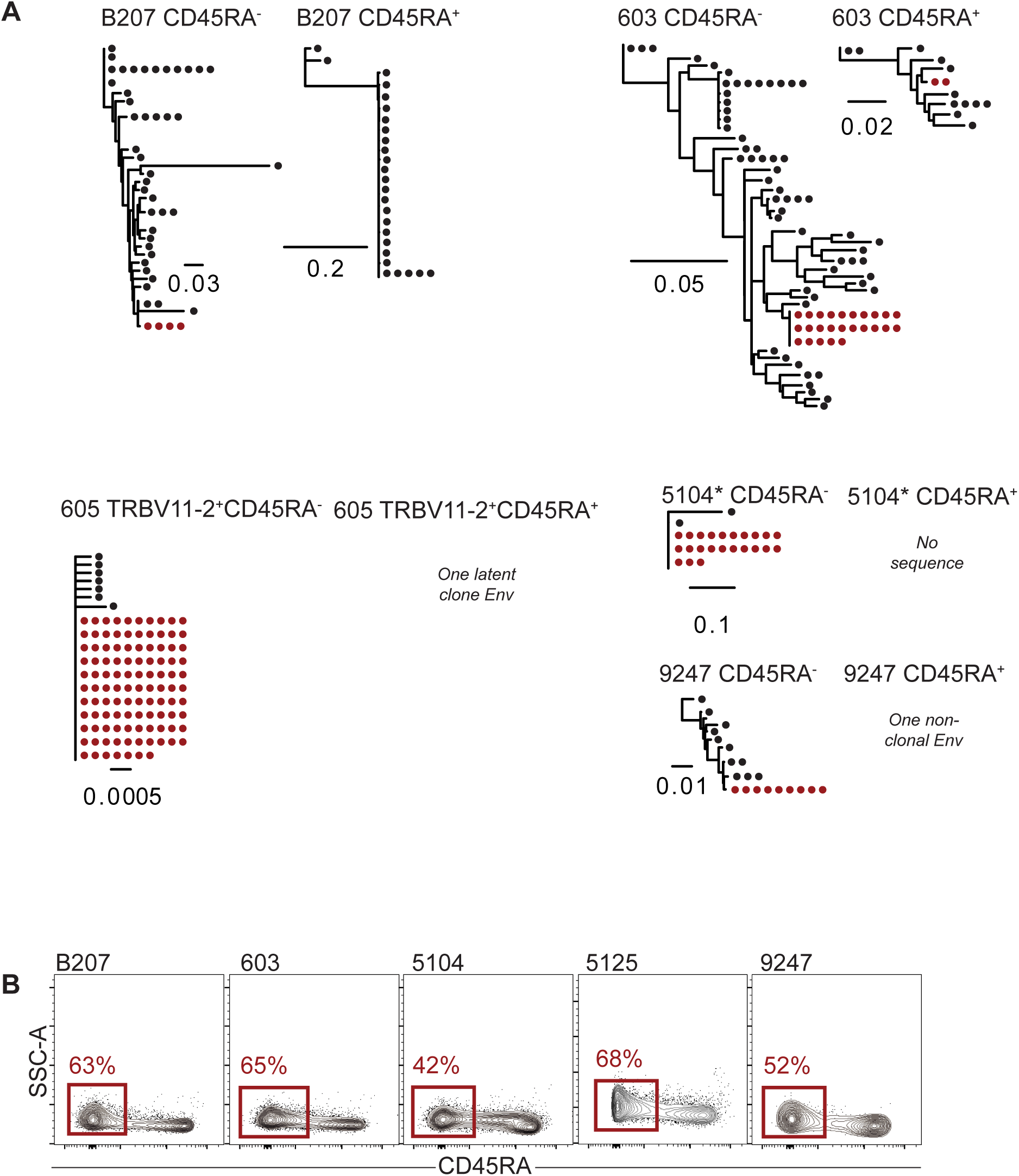
The latent clone of interest is enriched in the CD45RA^-^ compartment. Maximum-likelihood phylogenetic trees show *env* gene of the clone of interest marked in red. The scale bars indicate the number of substitutions per site. Asterisks indicate trees based on *env* sequences obtained from Q4PCR ^50^. (A) Magnetic negative selection of CD4^+^ CD45RA^-^ memory T cells enriches the latent clone of interest in all 6 individuals. (B) Flow cytometry plots indicating the fraction of CD45RA^-^ cells among CD4^+^ T cells.

**Figure S3.**
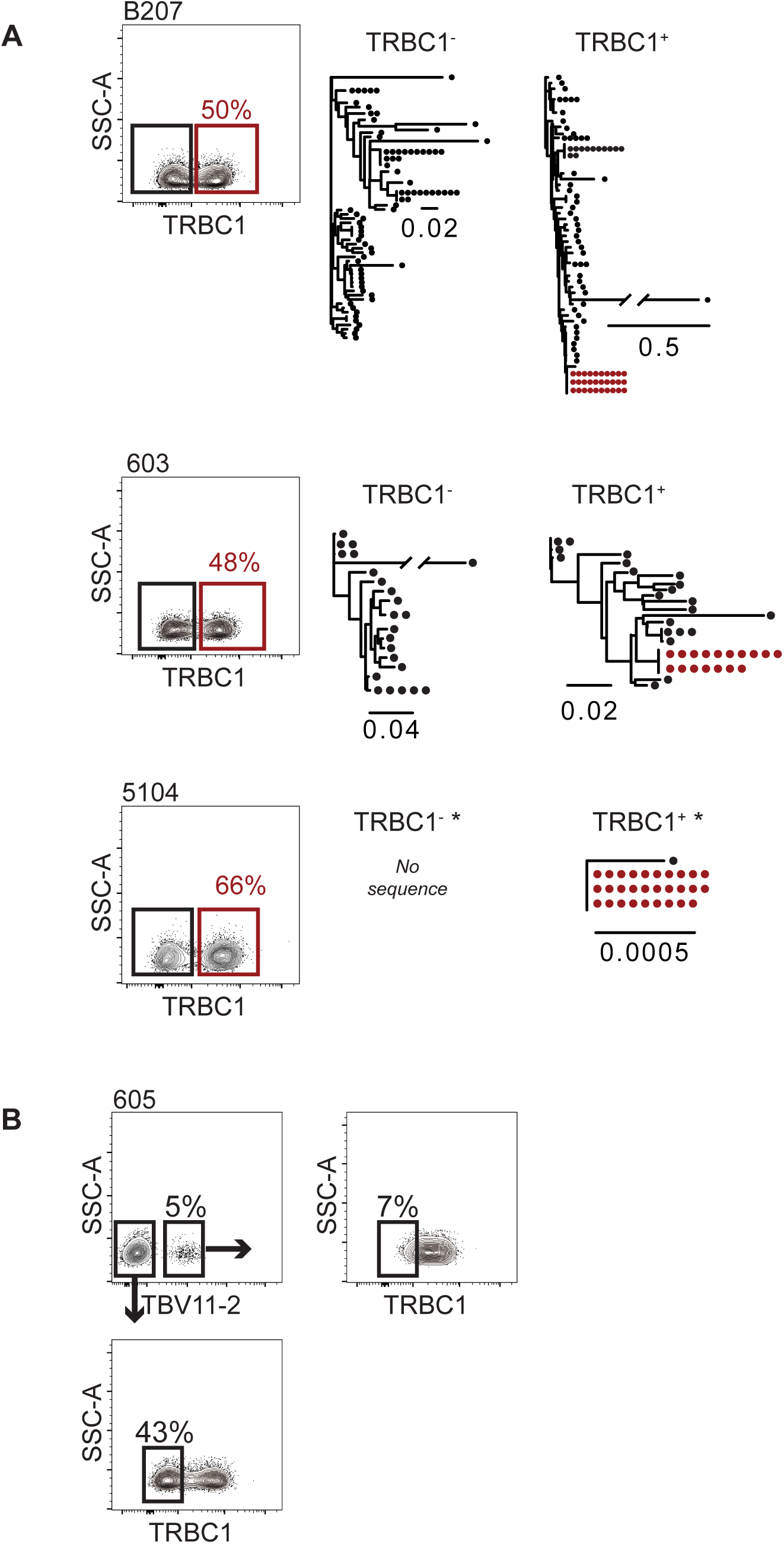
Identification of the TRBC domain that is expressed by the latent clone of interest. Red outlines indicate the population containing the clone of interest. Maximum-likelihood phylogenetic trees show *env* gene of the clone of interest marked in red. The scale bars indicate the number of substitutions per site. Asterisks indicate trees based on *env* sequences obtained from Q4PCR ^50^. (A) Flow cytometry plots show TRBC1 staining for individuals **B207**, **603**, and **5104**. The fraction of TRBC1^+^ CD4^+^ T cells is indicated. (B) Flow cytometry plots show TRBC1 staining for TRBV2^-^ and TRBV11-2^+^ cells in individual **605**. The fraction cells within a gate is indicated above the gate.

## Acknowledgments

We thank all study participants who devoted time to our research, the Rockefeller University Hospital Research support office and nursing staff, and all members of the M.C.N. laboratory for discussions, and M. Jankovic and G. Scrivanti for laboratory support. We thank C. Zhao, H. Duan, C. Lai, and S. Huang from the Genomics Resource Center at the Rockefeller University for preparing and sequencing the 10x Genomics libraries. We also thank K. Chhosphel and K. M. Gordon for operating the cell sorters.

This work was supported by the National Institutes of Health (grants 1U01AI129825 to M.C., UM1 AI100663 and R01AI129795 to M.C.N.); REACH Delaney (grant UM1 AI164565 to M.C.), the Einstein-Rockefeller-CUNY Center for AIDS Research (grant 1P30AI124414-01A1); BEAT-HIV Delaney (grant UM1 AI126620 to M.C.) and the Robertson Fund. L.B.C. is supported by the Delaney AIDS Research Enterprise (DARE) (1UM1AI126611-01), NIH REACH Martin Delaney Collaboratory (1UM1AI164565-01), and the Bill and Melinda Gates Foundation (INV-002707). C.G. was supported by the Robert S. Wennett postdoctoral fellowship, the Shapiro-Silverberg Fund for the Advancement of Translational Research and by the National Center for Advancing Translational Sciences (National Institutes of Health Clinical and Translational Science Award program, grant UL1 TR001866).

M.C.N. is a Howard Hughes Medical Institute (HHMI) Investigator. This article is subject to HHMI’s Open Access to Publications policy. HHMI lab heads have previously granted a nonexclusive CC BY 4.0 license to the public and a sublicensable license to HHMI in their research articles. Pursuant to those licenses, the author-accepted manuscript of this article can be made freely available under a CC BY 4.0 license immediately upon publication.

## Author contributions

G.H.J.W, Y.B-O., L.B.C., M.J., and M.C.N conceived and designed experiments. C.G. and M.C. recruited participants, supervised sample collection, and collected clinical data. G.H.J.W. performed the research. G.H.J.W., T.Y.O., V.R., L.B.C., M.J., and M.C.N analyzed data. T.Y.O. and V.R. performed bioinformatic analysis. H.H. and G.B. gave critical advice for cell sorting. G.H.J.W., L.B.C., M.J., and M.C.N. wrote the manuscript with help from all co-authors.

## Materials and methods

### Study design and participants

Study participants were recruited at the Rockefeller University hospital and gave informed written consent before participation in the studies. The study protocols and procedures met the standards of Good Clinical Practice and were approved by the institutional review board of the Rockefeller University. After leukapheresis, peripheral blood mononuclear cells (PBMCs) were isolated by Ficoll separation and stored in aliquots in liquid nitrogen. At the time of sample collection, all study participants were receiving ART and were aviremic.

### Magnetic-activated cell sorting of CD4 T cells and memory cells

All procedures were performed while maintaining cells at 4 °C. CD4^+^ T cells were negatively selected from PBMCs with a CD4^+^ T cell isolation kit (Miltenyi, cat. 130-096-533). For memory cell separation, CD4^+^ T cells were negatively selected using magnetic CD45RA MicroBeads (Miltenyi, cat. 130-045-901).

### Fluorescence-activated cell sorting

All procedures were performed while maintaining cells at 4 °C. Cells were incubated with Fc-blocking reagent (Miltenyi, cat. 130-059-901). Fixable Viability Dye eFluor 780 (Invitrogen, cat. 65-0865-14) was used for live/dead cell staining. The following antibodies were used for surface staining: PerCP/Cy5.5 anti-human CD4 (BioLegend, cat. 317428), PacificBlue anti-human CD3 (BioLegend, cat. 300431), Brilliant Violet 605 anti-human TCR Cβ1 (BD, cat. 747979), FITC anti-human TCR Vβ17 (Beckman Coulter, cat. IM1234), FITC anti-human TCR Vβ21.3 (Backman Coulter, cat. IM1483), FITC anti-human TCR Vβ7.2 (Beckman Coulter, cat. B06666), and Beta Mark TCR Vbeta Repertoire Kit (Beckman Coulter, cat. IM3497). The Vbeta Repertoire Kit contains 24 different anti-TRBV antibodies that come in 8 vials (A – H). In each vial, there are three antibodies conjugated with PE, or FITC, or PE FITC.

For enrichment, the following population was sorted after MACS for CD4^+^CD45RA^-^ T cells: individuals B207 and 5104 CD3^+^CD4^+^TRBC1^+^TRBV^-^ lymphocytes; individual 603 CD4^+^TRBC1^+^TRBV19^+^ lymphocytes; individual 605 CD4^+^TRBV11-2^+^ lymphocytes; individual 5125 CD4^+^TRBC1^+^TRBV2^+^ lymphocytes; individual 9247 CD4^+^TRBC1^-^TRBV4-3^+^ lymphocytes. Sorts were performed on BD FACS Aria IIu and BD FACS Aria III.

### Flowcytometric analysis

Flowcytometry data was analysed with FACSDiva software (version 2.0.2) and FlowJo version 10.8.1 (both by BD).

### Near-full length HIV provirus amplification by Q4 PCR and sequencing from bulk sorted cells

gDNA was extracted from sorted cells, quantified, and limiting dilution performed as described^26^. Near-full length genome amplification by Q4PCR was performed as described ^50^. PCR amplicons were sequenced with MiSeq 300 cycle v2 kits (Illumina, cat. MS-102-2002). Sequences were assembled with the Defective and Intact HIV Genome Assembler ^26^.

### *Env* amplification and sequencing from bulk sorted cells

gDNA was extracted from sorted cells, quantified, and limiting dilution performed as described^26^. All *env* amplifications were run with Platinum™ Taq DNA Polymerase High Fidelity (ThermoFisher, cat. 11304011) by nested PCR. For individual 605, the first PCR included the primers envB3out (5′-TTGCTACTTGTGATTGCTCCATGT–3′)^11^ and B3F3 (5 -′TGGAAAGGTGAAGGGGCAGT-AGTAATAC-3′) ^90^ at 94°C for 2min; (94°C for 15s, 60.4°C for 30s, and 68°C) for 6 min for 40 cycles; and 68°C for 15 min. For individuals B207, 603, 5104, 5125, and 9247, the first PCR as well as the second PCR for all individuals was performed as previously described ^11^. PCR amplicons were sequenced with MiSeq 300 cycle v2 kits (Illumina, cat. MS-102-2002). Sequences were assembled with the Defective and Intact HIV Genome Assembler ^26^.

### Phylogenetic trees

*Env* sequences were aligned to the HIV HXB2CG *env* sequence using Geneious Prime software (version 11.0.12, Biomatters). Maximum-likelihood phylogenetic trees were built with PHYML, substitution model HKY85, without bootstrapping, to identify identical *env* sequences by clustering.

### 10x Genomics

10x Genomics gene expression and V(D)J libraries were generated with the Chromium Single Cell 5’ Library & Gel Bead Kit (cat. PN-1000014) and Chromium Single Cell V(D)J Enrichment Kit, Human T Cell (cat. PN-1000005) as described in the 10x Genomics protocol. 5’ expression library was sequenced with NovaSeq 6000 S1 (100 cycles) (Illumina, cat. 20012865) and the V(D)J library was sequenced with NextSeq 500/550 Mid Output Kit v2.5 (300 Cycles) (Illumina, cat. 20024905).

### Latent clone TCR identification

Quiescent latent cells were enriched based on CD45RA, TRBC, and TRBV, and 5 cells per well were sorted into a 96-well plate containing TCL buffer (Qiagen, cat. 1031576) with 1 % beta- Mercaptethanol and snap frozen on dry ice. Plates were stored at -80 °C until further use. After thawing, magnetic bead clean-up was performed with Agencourt RNAClean XP magnetic beads (Beckman Coulter, cat. A63987). TCR mRNA was reverse transcribed into cDNA with SuperScript III Reverse Transcriptase (Invitrogen, cat. 18080044) and the following primers: AC1R (5’-ACACATCAGAATCCTTACTTTG-3’), BC1R (5’-CAGTATCTGGAGTCATTGA-3’) ^91^. Then, a second magnetic bead clean-up, binding TCR cDNA and gDNA, was performed, and *env* amplification from gDNA was performed for individuals 603, 5104, 5125, and 9247 as described above. Only for individual 605, a shortened nested PCR was performed to amplify a ∼ 500 bp amplicon of the *env* gene in which the expanded intact latent clone differed from other proviruses that were enriched. The first PCR was performed with primers Env 1,133 F1 (5’-GAGGGGAATTTTTCTACTGTAACAC-3’) and Env 1,956 R1 (5’- GTTCTGCGAATTTT-CAATTAAGGTG -3’) at 94°C for 2min; (94°C for 15s, 56°C for 30s, and 68°C for 1min) for 35 cycles; and 68°C for 15 min. The second PCR was performed with primers Env 1,240 F2 (5’- ATCACACTCCGATGC-AGAATAAAAC-3’) and Env 1,765 R2 (5’-TTAGGTATCT-TTCCACAGCCAGTAC-3’) at 94°C for 2min; 94°C for 15s, (58°C for 30s, and 68°C for 1min) for 35 cycles; and 68°C for 15 min.

TCR amplification and sequencing were performed with primers as previously described ^92^. TCR α and β chain cDNA was amplified and barcoded separately and in duplicates. All PCR products were combined at equal proportions, followed by magnetic bead clean up and run on a 1.5 % agarose gel. The band between 300 and 400 bp was excised, purified with NucleoSpin Gel and PCR Clean-up kit (Quiagen, cat. 740609.50) and sequenced on the MiSeq platform with a MiSeq 600 cycle v3 kit (Illumina, cat. MS-102-3003).

PhiX-derived reads were removed using from downstream analysis using a k-mer based approach implemented by bbduk.sh from BBTools v38.72 (https://sourceforge.net/projects/bbmap/). Samples were demultiplexed according to the combination of oligos used to uniquely identify the plate, row, column they were placed ^92^. The quality control check was performed with Trim Galore package v0.6.7 (https://github.com/FelixKrueger/TrimGalore ) to trim Illumina adapters and low-quality bases. Paired-overlapping reads were exported into a single read by BBMerge. Subsequently, TRUST4 ^93^ was used to reconstruct and annotate the T cell receptor (TCR) sequences. There is a chance of incorrect TCR assignment if the sequencing errors occur at the barcodes present in reads. A threshold was established based on the number of reads used to assembly a specific TCR contig assigned to an empty well. We obtained the putative clonotypes, defined by the 10x Genomics V(D)J single-cell sequencing, that shared either the α or β chain found in that well. The most frequent clonotype associated with Env^+^ wells in each individual was selected as the clonotype associated with proviral integration.

### Single-cell RNA-seq and single-cell TCR-seq processing

Single-cell RNA-seq binary base call (BCL) files were demultiplexed and converted into FASTQ files using BCLtoFastq prior to alignment to hg38 with CellRanger (v4.0.0) and analyzed in R studio with Seurat (v4). Cells with a mitochondrial proportion greater than 5 % and/or a feature count <200 or >2,500 were discarded. Sample batches were combined, normalized and scaled with SCTransform. Uniform Manifold Approximation and Projection (UMAP) clustering was performed selecting the first thirty principal components. Single-cell TCR-seq FASTQs were aligned with CellRanger (v4.0.0) to the default CellRanger VDJ reference. Output contig annotations were filtered and analyzed in R studio with Seurat.

### Mapping scRNA-seq to CD4^+^ T cells reference

CD4^+^ T cell population was extracted from published human peripheral blood cells multimodal annotated reference ^67^. The UMAP reference from extracted CD4^+^ T population was recreated using the first 50 principal components of the RNA expression slot and the cells from each individual were anchored and mapped utilizing the FindTransferAnchors and MapQuery functions from Seurat (reference.reduction = "pca", dims = 1:50, reduction.model = umap).

