## Supplementary material for "Distinct gene expression by expanded clones of quiescent memory CD4^+^ T cells harboring intact latent HIV-1 proviruses": Table S1

| **ID** | **Age** | **Gender** | **Race** | **Year HIV-1 Dx** | **Year ART initiation** | **Uninterr. ART (years)** | **Viral load (copies/mL)** | **CD4 T cell count** | **Reported nadir** | **ART regimen** |
| --- | --- | --- | --- | --- | --- | --- | --- | --- | --- | --- |
| **B207** | 47 | M | White/Hispanic | 10 | 10 | 9 | <20 | 724 | 50 | EFV/TDF/FTC |
| **603** | 43 | M | White/Hispanic | 12 | 10 | 10 | <20 | 300 | 693 | EFV/TDF/FTC |
| **605** | 36 | M | White/Hispanic | 15 | 14 | 2 | <20 | 372 | 524 | RPV/TDF/FTC |
| **5104** | 35 | M | Black | 7 | 7 | 7 | <20 | 400 | 606 | BIC/TAF/FTC |
| **5125** | 34 | M | Black | 11 | 10 | 2 | <20 | 450 | 1006 | DTG/TDF/FTC |
| **9247** | 31 | M | Black | 6 | 6 | 6 | <20 | 728 | 600 | EVG/cobi/TAF/FTC |

**Table S1: Clinical characteristics of study participants.**

Dx: diagnosis; Uninterr: uninterrupted; ART: antiretroviral treatment; EVG: elvitegravir; Cobi: cobicistat; TDF: tenofovir disoproxil fumarate; FTC: emtricitabine; RPV: rilpivirine; TAF: tenofovir alafenamide fumurate; BIC: bictegravir.
