## Supplementary material for "Distinct gene expression by expanded clones of quiescent memory CD4^+^ T cells harboring intact latent HIV-1 proviruses": Table S2

|  | **603** | **605** | **B207** | **5104** | **5125** | **9247** |
| --- | --- | --- | --- | --- | --- | --- |
| **CUPM Q4PCR** | ND | ND | ND | 210 | 28 | 15^1^ |
| **CUPM**  **Env PCR** | 13 | 431 | 164 | ND | ND | ND |
| **Latent clone inducible *in vitro*** | Yes^2^ | Yes^2^ | Yes^2,3^ | Yes^3^ | ND | No |
| **Integration site** | ND | ND | ZPF30^3^ | ATP2B4, DUXL6^3^ | ND | ND |

**Table S2: Frequency, inducibility, and integration site of the clone of interest in CD4^+^ T cells**

CUPM: Clonal units of clone of interest per million CD4^+^ T cells

1	Gaebler, C. *et al.* Sequence Evaluation and Comparative Analysis of Novel Assays for Intact Proviral HIV-1 DNA. *J Virol* **95**, doi:10.1128/JVI.01986-20 (2021).

2	Cohn, L. B. *et al.* Clonal CD4(+) T cells in the HIV-1 latent reservoir display a distinct gene profile upon reactivation. *Nat Med* **24**, 604-609, doi:10.1038/s41591-018-0017-7 (2018).

3	Huang, A. S. *et al.* Integration features of intact latent HIV-1 in CD4+ T cell clones contribute to viral persistence. *J Exp Med* **218**, doi:10.1084/jem.20211427 (2021).
