## Supplementary material for "Distinct gene expression by expanded clones of quiescent memory CD4^+^ T cells harboring intact latent HIV-1 proviruses": Table S3

|  | **CD45RA** | **TRBC** | **TRBV** | **combined** |
| --- | --- | --- | --- | --- |
| **603** | 1.5 | 2.2 | 14.3 | 47.2 |
| **605** | ~2 (estimate) | - | 20 | 40 |
| **B207** | 1.6 | 2 | 2.9 | 9.3 |
| **5104** | 2.4 | 1.6 | 2.9 | 11.1 |
| **5125** | 1.5 | 1.8 | 20 | 54 |
| **9247** | 1.9 | 5 | 71 | 674.5 |

**Table S3: Relative enrichment per marker based on flow cytometry.**
