## Supplementary material for "Distinct gene expression by expanded clones of quiescent memory CD4^+^ T cells harboring intact latent HIV-1 proviruses": Table S4

|  | **Latent clone *env* copies per**  **10^4^ CD4^+^ T cells** | **Specific TCR per 10^4^ CD4^+^ T cells** | **Ratio TCR/*env*** |
| --- | --- | --- | --- |
| **603** | 20 | 45 | 2.3 |
| **605** | 163 | 502 | 3.1 |
| **B207** | 6 | 14 | 2.3 |
| **5104** | 14 | 16 | 1.1 |
| **5125** | 15 | 17 | 1.1 |
| **9247** | 18 | 23 | 1.3 |

**Table S4: Comparison between *env* copies per 10^4^ CD4^+^ T cells (based on *env* PCR) and the frequency of the latent clone TCR (based on 10x Genomics TCR sequencing) after enrichment based on CD45RA, TRBV, and TRBC.**
