## Supplementary material for "Distinct gene expression by expanded clones of quiescent memory CD4^+^ T cells harboring intact latent HIV-1 proviruses": Table S5

| **Number of cells analyzed by 10x genomics single cell gene expression** | | | | | | | |
| --- | --- | --- | --- | --- | --- | --- | --- |
| **Individual** | **603** | **605** | **B207** | **5104** | **5125** | **9247** | **Sum** |
| **# of cells** | 16,224 | 17,357 | 27,472 | 21,143 | 12,173 | 14,848 | 109,217 |

| **Individual** | **603** | | | **605** | | | **B207** | | | **5104** | | | **5125** | | | **9247** | | |
| --- | --- | --- | --- | --- | --- | --- | --- | --- | --- | --- | --- | --- | --- | --- | --- | --- | --- | --- |
| **Clone** | smaller (n=64) | latent (n=75) | bigger (n=84) | smaller (n=330) | latent (n=844) | bigger | smaller (n=40) | latent (n=41) | bigger (n=42) | smaller (n=32) | latent (n=35) | bigger (n=38) | smaller (n=21) | latent (n=21) | bigger (n=22) | smaller (n=34) | latent (n=34) | bigger (n=35) |
| **Cluster 0** | 17.2 | 18.7 | 34.5 | 8.5 | 7.8 | - | 0.0 | 12.2 | 35.7 | 43.8 | 17.1 | 0.0 | 33.3 | 9.5 | 9.1 | 8.8 | 0.0 | 0.0 |
| **Cluster 1** | 54.7 | 0.0 | 4.8 | 68.2 | 2.7 | - | 5.0 | 0.0 | 26.2 | 0.0 | 5.7 | 0.0 | 19.0 | 14.3 | 36.4 | 2.9 | 2.9 | 5.7 |
| **Cluster 2** | 1.6 | 0.0 | 0.0 | 0.0 | 0.0 | - | 0.0 | 0.0 | 11.9 | 40.6 | 8.6 | 0.0 | 28.6 | 4.8 | 27.3 | 20.6 | 23.5 | 34.3 |
| **Cluster 3** | 1.6 | 0.0 | 0.0 | 1.5 | 0.6 | - | 5.0 | 4.9 | 4.8 | 3.1 | 0.0 | 0.0 | 9.5 | 0.0 | 9.1 | 0.0 | 0.0 | 0.0 |
| **Cluster 4** | 3.1 | 2.7 | 7.1 | 4.5 | 1.2 | - | 25.0 | 2.4 | 0.0 | 3.1 | 2.9 | 0.0 | 0.0 | 4.8 | 13.6 | 35.3 | 8.8 | 31.4 |
| **Cluster 5** | 4.7 | 10.7 | 10.7 | 11.5 | 1.7 | - | 0.0 | 0.0 | 4.8 | 6.3 | 0.0 | 0.0 | 4.8 | 4.8 | 0.0 | 5.9 | 0.0 | 5.7 |
| **Cluster 6** | 4.7 | 0.0 | 0.0 | 0.0 | 0.7 | - | 0.0 | 0.0 | 2.4 | 0.0 | 0.0 | 0.0 | 4.8 | 0.0 | 4.5 | 23.5 | 35.3 | 22.9 |
| **Cluster 7** | 0.0 | 60.0 | 23.8 | 2.1 | 63.7 | - | 0.0 | 73.2 | 4.8 | 3.1 | 65.7 | 0.0 | 0.0 | 47.6 | 0.0 | 0.0 | 29.4 | 0.0 |
| **Cluster 8** | 0.0 | 8.0 | 0.0 | 0.0 | 3.8 | - | 0.0 | 7.3 | 0.0 | 0.0 | 0.0 | 100.0 | 0.0 | 9.5 | 0.0 | 0.0 | 0.0 | 0.0 |
| **Cluster 9** | 0.0 | 0.0 | 0.0 | 0.0 | 0.5 | - | 2.5 | 0.0 | 0.0 | 0.0 | 0.0 | 0.0 | 0.0 | 0.0 | 0.0 | 0.0 | 0.0 | 0.0 |
| **Cluster 10** | 0.0 | 0.0 | 0.0 | 0.3 | 0.2 | - | 62.5 | 0.0 | 0.0 | 0.0 | 0.0 | 0.0 | 0.0 | 0.0 | 0.0 | 0.0 | 0.0 | 0.0 |
| **Cluster 11** | 7.8 | 0.0 | 16.7 | 1.8 | 16.1 | - | 0.0 | 0.0 | 9.5 | 0.0 | 0.0 | 0.0 | 0.0 | 4.8 | 0.0 | 0.0 | 0.0 | 0.0 |
| **Cluster 12** | 0.0 | 0.0 | 2.4 | 0.0 | 0.4 | - | 0.0 | 0.0 | 0.0 | 0.0 | 0.0 | 0.0 | 0.0 | 0.0 | 0.0 | 2.9 | 0.0 | 0.0 |
| **Cluster 13** | 4.7 | 0.0 | 0.0 | 0.6 | 0.2 | - | 0.0 | 0.0 | 0.0 | 0.0 | 0.0 | 0.0 | 0.0 | 0.0 | 0.0 | 0.0 | 0.0 | 0.0 |
| **Cluster 14** | 0.0 | 0.0 | 0.0 | 0.9 | 0.4 | - | 0.0 | 0.0 | 0.0 | 0.0 | 0.0 | 0.0 | 0.0 | 0.0 | 0.0 | 0.0 | 0.0 | 0.0 |
| **Sum** | 100 | 100 | 100 | 100 | 100 | - | 100 | 100 | 100 | 100 | 100 | 100 | 100 | 100 | 100 | 100 | 100 | 100 |

**Table S5: Upper Distribution [%] of the latent clone, the next smaller, and the next bigger clone per individual over the 15 gene expression clusters**
