## Supplementary material for "Distinct gene expression by expanded clones of quiescent memory CD4^+^ T cells harboring intact latent HIV-1 proviruses": Table S6

| **Ensembl gene ID** | **External gene name** | **p value** | **Average log2 fold change** |
| --- | --- | --- | --- |
| ENSG00000115687 | PASK | 2.5074E-273 | -0.937004582 |
| ENSG00000188404 | SELL | 1.1432E-303 | -0.757375402 |
| ENSG00000168209 | DDIT4 | 4.2236E-135 | -0.753863566 |
| ENSG00000070756 | PABPC1 | 1.3615E-275 | -0.577338107 |
| ENSG00000074800 | ENO1 | 3.9911E-72 | -0.552495574 |
| ENSG00000273149 | AL138963.4 | 1.69448E-43 | -0.503468146 |
| ENSG00000111669 | TPI1 | 3.75841E-56 | -0.502939955 |
| ENSG00000095794 | CREM | 7.5629E-48 | -0.48733596 |
| ENSG00000157601 | MX1 | 3.44639E-38 | -0.463685933 |
| ENSG00000149212 | SESN3 | 2.09806E-96 | -0.458160858 |
| ENSG00000126353 | CCR7 | 5.6185E-113 | -0.438022627 |
| ENSG00000177410 | ZFAS1 | 9.1716E-145 | -0.432744809 |
| ENSG00000173762 | CD7 | 1.2733E-119 | -0.422179842 |
| ENSG00000059804 | SLC2A3 | 4.68348E-68 | -0.401753273 |
| ENSG00000197061 | HIST1H4C | 1.74666E-54 | -0.367052221 |
| ENSG00000234741 | GAS5 | 1.8049E-124 | -0.365481063 |
| ENSG00000134333 | LDHA | 3.62429E-18 | -0.356852098 |
| ENSG00000197989 | SNHG12 | 2.6524E-59 | -0.354230806 |
| ENSG00000105220 | GPI | 2.30295E-38 | -0.33651571 |
| ENSG00000146278 | PNRC1 | 2.56242E-66 | -0.33232545 |
| ENSG00000196352 | CD55 | 3.58249E-86 | -0.331613182 |
| ENSG00000269028 | MTRNR2L12 | 9.28887E-42 | -0.326076812 |
| ENSG00000251562 | MALAT1 | 4.23243E-61 | -0.326073295 |
| ENSG00000240972 | MIF | 1.16951E-48 | -0.325935464 |
| ENSG00000067225 | PKM | 3.0003E-53 | -0.324450491 |
| ENSG00000166012 | TAF1D | 4.1997E-62 | -0.323635813 |
| ENSG00000130066 | SAT1 | 6.88644E-24 | -0.320711104 |
| ENSG00000161011 | SQSTM1 | 3.25091E-23 | -0.307785538 |
| ENSG00000181163 | NPM1 | 1.7052E-145 | -0.295019474 |
| ENSG00000081059 | TCF7 | 2.53957E-59 | -0.293559954 |
| ENSG00000100906 | NFKBIA | 6.90414E-22 | -0.286544704 |
| ENSG00000138795 | LEF1 | 4.65739E-63 | -0.283635236 |
| ENSG00000102144 | PGK1 | 3.47758E-18 | -0.283588302 |
| ENSG00000104765 | BNIP3L | 4.68824E-42 | -0.280401542 |
| ENSG00000144381 | HSPD1 | 1.33312E-51 | -0.27754675 |
| ENSG00000096384 | HSP90AB1 | 6.19504E-51 | -0.27513593 |
| ENSG00000114023 | FAM162A | 3.98958E-38 | -0.271026449 |
| ENSG00000167658 | EEF2 | 2.0542E-116 | -0.263184273 |
| ENSG00000081320 | STK17B | 1.01417E-40 | -0.263109056 |
| ENSG00000105193 | RPS16 | 7.12703E-82 | -0.25357902 |
| ENSG00000213145 | CRIP1 | 1.84831E-74 | 0.251381711 |
| ENSG00000204642 | HLA-F | 1.67539E-32 | 0.252987681 |
| ENSG00000064666 | CNN2 | 7.14478E-34 | 0.254603703 |
| ENSG00000136167 | LCP1 | 4.88255E-38 | 0.255115632 |
| ENSG00000145247 | OCIAD2 | 1.35036E-33 | 0.255136956 |
| ENSG00000135441 | BLOC1S1 | 8.55057E-29 | 0.256708541 |
| ENSG00000217555 | CKLF | 6.26258E-37 | 0.268552743 |
| ENSG00000179218 | CALR | 5.15079E-26 | 0.269118004 |
| ENSG00000166710 | B2M | 0 | 0.270596986 |
| ENSG00000034713 | GABARAPL2 | 7.97742E-47 | 0.271789708 |
| ENSG00000213626 | LBH | 1.70427E-35 | 0.275234928 |
| ENSG00000158062 | UBXN11 | 2.96782E-31 | 0.277861974 |
| ENSG00000135046 | ANXA1 | 6.7754E-104 | 0.277916253 |
| ENSG00000240065 | PSMB9 | 9.1741E-63 | 0.278633877 |
| ENSG00000165929 | TC2N | 3.38931E-34 | 0.282431558 |
| ENSG00000136810 | TXN | 4.66479E-33 | 0.284410494 |
| ENSG00000115232 | ITGA4 | 4.07453E-43 | 0.296034223 |
| ENSG00000234745 | HLA-B | 2.442E-301 | 0.299434559 |
| ENSG00000182718 | ANXA2 | 1.2129E-48 | 0.300086815 |
| ENSG00000105404 | RABAC1 | 3.41798E-59 | 0.300856113 |
| ENSG00000108518 | PFN1 | 5.1347E-192 | 0.308425928 |
| ENSG00000075624 | ACTB | 3.1245E-173 | 0.312823868 |
| ENSG00000163191 | S100A11 | 5.8168E-123 | 0.313585328 |
| ENSG00000213719 | CLIC1 | 3.2501E-80 | 0.321146717 |
| ENSG00000126246 | IGFLR1 | 1.12795E-32 | 0.321234367 |
| ENSG00000027869 | SH2D2A | 1.2519E-59 | 0.325040074 |
| ENSG00000008517 | IL32 | 1.885E-203 | 0.326877861 |
| ENSG00000100300 | TSPO | 4.23815E-58 | 0.328869744 |
| ENSG00000130592 | LSP1 | 5.13406E-85 | 0.335795569 |
| ENSG00000170571 | EMB | 1.30243E-48 | 0.340520299 |
| ENSG00000110324 | IL10RA | 7.30307E-71 | 0.345878196 |
| ENSG00000092841 | MYL6 | 1.8186E-212 | 0.349021829 |
| ENSG00000160255 | ITGB2 | 1.63101E-55 | 0.376681806 |
| ENSG00000197747 | S100A10 | 5.2906E-220 | 0.381136641 |
| ENSG00000169442 | CD52 | 2.9441E-231 | 0.386540002 |
| ENSG00000132965 | ALOX5AP | 2.30411E-89 | 0.390330172 |
| ENSG00000148362 | PAXX | 1.9717E-94 | 0.391613282 |
| ENSG00000197540 | GZMM | 5.469E-80 | 0.411312119 |
| ENSG00000002586 | CD99 | 1.1703E-191 | 0.411377898 |
| ENSG00000111796 | KLRB1 | 8.73173E-56 | 0.434616046 |
| ENSG00000196154 | S100A4 | 9.403E-205 | 0.446707794 |
| ENSG00000142669 | SH3BGRL3 | 0 | 0.473220735 |
| ENSG00000126264 | HCST | 3.4505E-135 | 0.484457968 |
| ENSG00000103187 | COTL1 | 8.1033E-136 | 0.491151274 |
| ENSG00000100097 | LGALS1 | 3.0001E-118 | 0.520481509 |
| ENSG00000116824 | CD2 | 7.0921E-219 | 0.527921045 |
| ENSG00000051523 | CYBA | 0 | 0.535424541 |
| ENSG00000133321 | PLAAT4 | 1.0403E-293 | 0.6130956 |
| ENSG00000235576 | LINC01871 | 6.8684E-252 | 0.712737658 |
| ENSG00000186810 | CXCR3 | 0 | 0.863806843 |
| ENSG00000145220 | LYAR | 0 | 0.91690397 |
| ENSG00000223865 | HLA-DPB1 | 0 | 1.005834612 |
| ENSG00000077984 | CST7 | 0 | 1.00668931 |
| ENSG00000231389 | HLA-DPA1 | 0 | 1.0517126 |
| ENSG00000145649 | GZMA | 0 | 1.159266639 |
| ENSG00000196126 | HLA-DRB1 | 0 | 1.189048765 |
| ENSG00000158050 | DUSP2 | 0 | 1.377812382 |
| ENSG00000271503 | CCL5 | 0 | 1.450021207 |
| ENSG00000019582 | CD74 | 0 | 1.478166065 |
| ENSG00000113088 | GZMK | 0 | 2.055910352 |

**Table S6: Differentially expressed genes (average log2fold < -2.5; average log2fold > 2.5) in cluster 7 compared to all other cluster**
