## Supplementary material for "Distinct gene expression by expanded clones of quiescent memory CD4^+^ T cells harboring intact latent HIV-1 proviruses": Table S7

|  | **603** | **605** | **B207** | **5104** | **5125** | **9247** |
| --- | --- | --- | --- | --- | --- | --- |
| **CD4 CTL** | 1.6% | 58.5% | 0.0% | 0.0% | 0.9% | 0.0% |
| **CD4 Naïve** | 0.0% | 1.0% | 0.0% | 0.0% | 0.0% | 0.0% |
| **CD4 Proliferating** | 0.0% | 0.0% | 0.0% | 0.0% | 0.0% | 0.0% |
| **CD4 TCM** | 0.1% | 2.2% | 0.0% | 0.1% | 0.1% | 0.2% |
| **CD4 TEM** | 6.1% | 21.9% | 2.4% | 1.1% | 3.8% | 2.0% |
| **Treg** | 0.0% | 0.4% | 0.0% | 0.0% | 0.0% | 0.0% |

**Table S7: Percentage of cell pertaining to the latent clone relative to cluster size**
