## Supplementary material for "Distinct gene expression by expanded clones of quiescent memory CD4^+^ T cells harboring intact latent HIV-1 proviruses": Table S8

| **Individual** | **Sample** | **Cells assayed** | **Sequences of clone of interest** | **Sequences of clone of interest per 10^6^ CD4^+^ T cells** | **Assay (Ref.)** | **doi (if applicable)** |
| --- | --- | --- | --- | --- | --- | --- |
| **603** | CD4^+^ | 1,228,800 | 16 | 13.0 | Env PCR |  |
|  | enriched | 49,920 | 102 | 2,043.0 | Env PCR |  |
| **605** | CD4^+^ | 576,000 | 248 | 430.5 | Env PCR |  |
|  | enriched | 3,072 | 50 | 16,276.0 | Env PCR |  |
| **B207** | CD4^+^ | 249,600 | 41 | 164.3 | Env PCR |  |
|  | enriched | 115,200 | 71 | 616.3 | Env PCR |  |
| **5104** | CD4^+^ | 576,000 | 121 | 210.0 | Q4PCR (Gaebler, submitted |  |
|  | enriched | 115,200 | 159 | 1,380.2 | Env PCR |  |
| **5125** | CD4^+^ | 1,440,000 | 40 | 27.8 | Q4PCR (Gaebler, submitted) |  |
|  | enriched | 64,512 | 95 | 1,472.6 | Env PCR |  |
| **9247** | CD4^+^ | 960,000 | 14 | 14.6 | Q4PCR (Gaebler et al., 2021) | https://doi.org/10.1128/JVI.01986-20 |
|  | enriched | 19,200 | 34 | 1,770.8 | Env PCR |  |

**Table S8: Number of cells analyzed and assay type before and after enrichment**
